# Metabolomics identifies endocannabinoid system remodeling during obesity associated inflammation in adipose tissue

**DOI:** 10.64898/2026.09.25.753352

**Authors:** Nasim Bararpour, Tiziana Caputo, Tatjana Sajic, Carine Winkler, Nicolas Guex, Béatrice Desvergne, Federica Gilardi, Aurélien Thomas

## Abstract

Visceral white adipose tissue (vAT) is more susceptible than subcutaneous white adipose tissue (scAT) to obesity-associated metaflammation, but the metabolic pathways accompanying this depot-specific transition remain incompletely defined. We profiled vAT and scAT from mice fed a control diet or a 60%-kcal high-fat diet (HFD) for 1 or 8 weeks. Untargeted metabolomics was integrated with targeted endocannabinoid quantification, gene-expression and promoter-associated chromatin analyses, adipocyte and stromal vascular fraction (SVF) measurements, cross-omics correlation analysis, and cell-culture perturbations. After 1 week, HFD induced relatively few metabolic changes, although vAT showed early alterations associated with sphingomyelin metabolism. By week 8, both depots exhibited extensive but distinct metabolic remodeling. The KEGG retrograde endocannabinoid-signaling pathway was the highest-ranked enriched pathway in vAT, with 2-arachidonoylglycerol (2-AG) among the most strongly altered metabolites. Targeted analysis confirmed selective 2-AG accumulation in vAT, accompanied by coordinated changes in genes and promoter-associated chromatin marks related to endocannabinoid synthesis, degradation and receptor expression. Cellular fractionation localized excess 2-AG predominantly to vAT adipocytes, whereas *Cnr2* was enriched in the SVF and induced by HFD specifically in vAT. Cross-omics integration further revealed a denser and predominantly inverse transcript–metabolite association network in vAT than in scAT. Using macrophage–adipocyte co-culture and secretome profiling, we demonstrated that cannabinoid stimulation shifts activated macrophage outputs away from pro-inflammatory cytokines (*Il1b*, *Ccl2*) toward extracellular matrix (ECM) proteoglycan remodeling and intercellular signaling.

**Graphical Abstract:** Early lipid remodeling in visceral adipose tissue is followed by coordinated endocannabinoid-system stimulation during sustained high-fat-diet feeding inducing inflammation

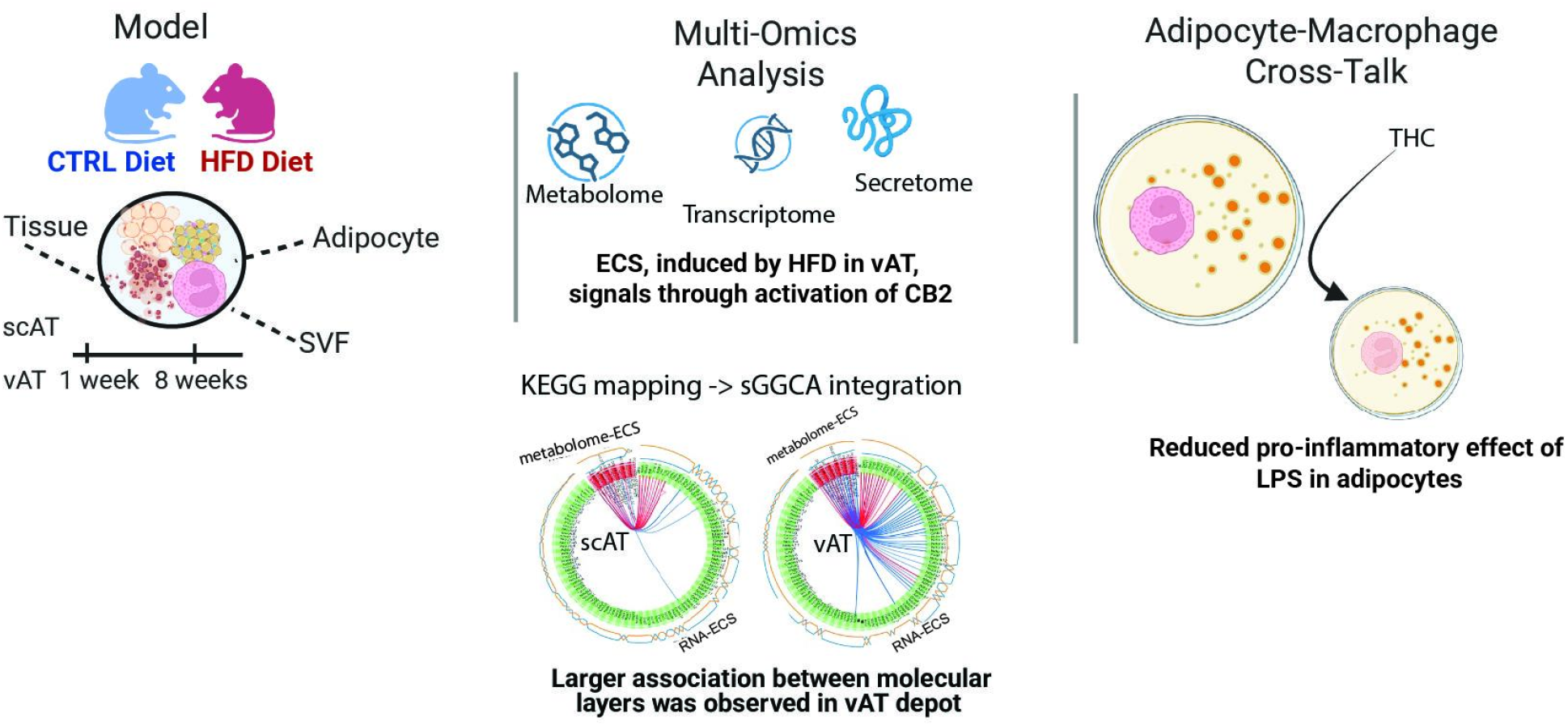

- Metabolic alterations were detectable after 1 week of HFD feeding in VAT.
- Sustained HFD exposure preferentially remodels the visceral adipose ECS, producing extensive and predominantly coordination between ECS-related transcriptional and metabolic profiles.
- In presence of both macrophage and adipocytes THC dampens the pro-inflammatory effect of LPS in adipocytes

## Introduction

The prevalence of obesity has increased markedly worldwide over the past three decades. Obesity is characterized by expansion of adipose tissue, an organ specialized in energy storage and metabolic regulation. Although increased adiposity does not invariably result in metabolic disease, obesity is frequently accompanied by chronic, low-grade systemic inflammation that contributes to insulin resistance and related cardiometabolic complications ^1^. White adipose tissue is an important early site of this inflammatory response, particularly the visceral depot surrounding the intra-abdominal organs. Compared with visceral adipose tissue (vAT), subcutaneous adipose tissue (scAT) is generally less susceptible to obesity-associated inflammation and metabolic dysfunction ^2^. Systematic comparison of these depots can therefore reveal molecular events associated with the transition from adaptive adipose expansion to tissue inflammation. We previously showed that mouse vAT and scAT differ in their ability to recruit and differentiate new adipocytes during high-fat-diet (HFD) feeding, contributing to their distinct susceptibility to inflammation ^3^.

During overnutrition, white adipose tissue undergoes extensive remodeling through adipocyte hypertrophy and, depending on the depot, the formation of new adipocytes. When tissue expansion exceeds its adaptive capacity, it is accompanied by oxidative and endoplasmic-reticulum stress, altered adipokine and cytokine secretion, extracellular-matrix remodeling and immune-cell recruitment. Adipose production of inflammatory mediators, including tumour necrosis factor-α, contributes to the development of obesity-associated insulin resistance ^4^. These cellular changes are accompanied by coordinated regulation of metabolic and signaling pathways, but the lipid-derived signals that distinguish visceral from subcutaneous adipose tissue during the early inflammatory transition remain incompletely understood.

The endocannabinoid system (ECS) is a lipid-signaling network involved in energy balance, adipose metabolism and immune regulation. Initially characterized in the central nervous system for its role in retrograde synaptic signaling, the ECS is also active in peripheral metabolic tissues, including adipose tissue. Its best-characterized endogenous ligands are 2-arachidonoylglycerol (2-AG) and N-arachidonoylethanolamine, or anandamide (AEA), which activate cannabinoid receptor type 1 (CB1) and type 2 (CB2) with different affinities and efficacies^5,6^.

Local ECS activity is controlled by ligand synthesis and degradation, precursor availability, receptor expression and cellular context. 2-AG is synthesized from diacylglycerol by the α and β isoforms of diacylglycerol lipase, *DAGLα* and *DAGLβ. DAGLα* is prominent in the nervous system, whereas *DAGLβ* has important functions in peripheral and immune-cell contexts. AEA can be generated from N-acyl-phosphatidylethanolamine through several enzymatic routes, including hydrolysis by N-acyl-phosphatidylethanolamine-specific phospholipase D. 2-AG is degraded primarily by monoacylglycerol lipase, with additional contributions from *ABHD6* and *ABHD12*, whereas AEA is hydrolysed mainly by fatty acid amide hydrolase. Endocannabinoid concentrations therefore reflect the balance among multiple synthetic and degradative pathways rather than the expression of a single enzyme.

Peripheral ECS activity is frequently altered in obesity, and circulating 2-AG has been associated particularly with visceral adiposity and adverse metabolic parameters^7,8^. Nevertheless, the net consequences of ECS remodeling within adipose tissue remain difficult to predict because they depend on ligand identity, receptor distribution, cellular composition and inflammatory state. CB1 signaling in adipocytes promotes lipid storage and may impair mitochondrial biogenesis, whereas CB2 is enriched in immune-cell populations and has shown context-dependent inflammatory effects. ECS remodeling could therefore have distinct, and potentially opposing, consequences in adipocytes and immune cells.

Metabolomics provides a direct means of measuring endocannabinoids, their lipid precursors and the broader metabolic environment integrating dietary exposure, enzyme activity and tissue state. We used metabolomics to compare vAT and scAT response following acute and sustained HFD feeding^3^. We revealed ECS-centered remodeling through untargeted metabolomics, absolute endocannabinoid quantification, integration with transcriptomic and promoter-associated chromatin profiles, analysis of separated adipocyte and stromal vascular fractions, cannabinoid perturbation of macrophage and adipocyte models, macrophage–adipocyte co-culture, and proteomic characterization of the macrophage secretome.

## Results

### Sustained high-fat feeding reveals divergent metabolic responses in visceral and subcutaneous adipose tissue

The experimental design, sampling time points and molecular profiling platforms are summarized in Figure 1. As expected, mice fed a high-fat diet (HFD) gained more weight than control-diet mice (Figure S1A). AT inflammatory response in these mice was previously characterized ^3^ DOI: 10.1038/s41366-023-01450-x(Caputo et al 2021, Sajic et al 2024). By week 8, vAT had reached its greatest expansion during the study and exhibited more pronounced macrophage infiltration and inflammatory-marker expression than scAT (Figure S1B) [1]. Phenotypic clustering based on body weight, circulating insulin, leptin and resistin, together with adipose expression of *Ccl2, Itgax* and *Cxcl12*, separated control mice from most HFD-fed animals^3^. A subset of HFD-fed mice remained phenotypically close to controls and was classified as low inflammation (Low-INFL HFD) because it exhibited limited vAT inflammation ^3,9^ (Figure S1C; see Method). Untargeted metabolomics was performed in both adipose depots using samples from 116 mice included in the study. Signal drift and batch effects were corrected using the DBnorm workflow^10^ (Figure S1D). Differential analysis indicated a limited metabolic response after 1 week of HFD feeding (Figure S2A). Significance criteria of Benjamini–Hochberg-adjusted(BH) P < 0.05 and |log2 fold change (log2FC)| > 1 have been applied for feature selection (see Method). Both depots showed early changes in metabolites associated with choline and glycerophospholipid metabolism, whereas vAT additionally displayed alterations related to sphingomyelin metabolism (Figure S2B–C). These early changes suggest that lipid remodeling, in vAT, precedes the pronounced inflammatory phenotype observed after sustained HFD exposure^11^.By week 8, the number of annotated metabolites and lipids meeting with predefined significance criteria, BH-adjusted P < 0.05 and log2FC> 1, had increased markedly (Figure S2D; Table S1A). Using the 8-week scAT response as an anchor, significantly altered features were followed across the other diet, time and depot contrasts (Figure S3A). Features detected selectively in 8-week scAT, including several unsaturated fatty acids, retinoic acid and selected phospholipids, further indicated a depot-specific component of the response (Table S1B). Nearly half of this response was shared with vAT at week 8. These shared alterations were not significant after 1 week of HFD feeding or between the 1- and 8-week control-diet groups, supporting an association with sustained HFD exposure rather than time alone. comparisons. Prominent shared signatures included reductions in lysophosphatidylcholines and unsaturated fatty acids, together with increases in ceramides, sphingomyelins and acylcarnitines (Figure S3B–C).

**Figure 1.**
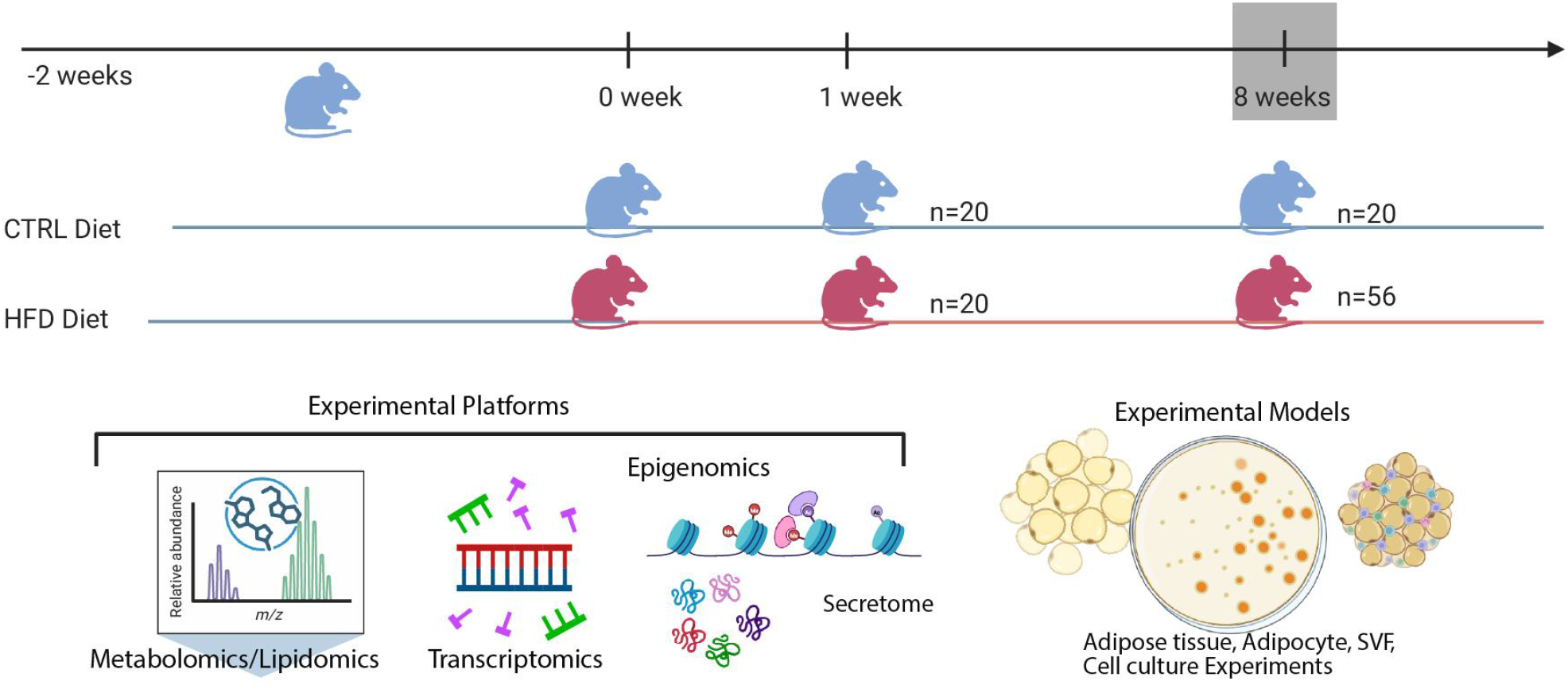
Study design and data acquisition platforms. Top panel: Four-week-old mice were fed a control chow diet for 2 weeks. At 6 weeks of age, mice were either switched to a high-fat diet (HFD) containing 60% fat or maintained on the control diet for 1 or 8 weeks. Visceral and subcutaneous adipose tissue (vAT and scAT) were collected at sacrifice for further analyses. Bottom panel: Multi-omics profiling was performed using untargeted metabolomics for discovery, followed by targeted metabolomics for validation of the findings. Data were combined with epigenomic profiling to assess chromatin accessibility and enhancer-associated histone modifications, including histone H3 lysine 27 acetylation (H3K27ac) and H3 lysine 4 monomethylation (H3K4me1) and transcriptomics data previously obtained (Caputo et al. 2021). Secretome profiling was performed in macrophage-adipocytes co-culture experiments.

Sustained HFD exposure (8-week) was the primary driver of the metabolic response in vAT(Figure S4A-C). Only a small proportion overlapped with the 1-week diet contrasts or with time-dependent changes in control-fed mice (Figure S4A; Table S1A). Thus, most of the vAT signature emerged between weeks 1 and 8 and was primarily associated with prolonged HFD exposure rather than age alone.

The 8-week vAT response was characterized predominantly by increased lipid species, including glycerophospholipids, sphingolipids and mono- or diacylglycerols (Figure S4B; Table S1C).

Increased ceramides, acylcarnitines and diacylglycerols were consistent with the hyperinsulinemia and pronounced inflammatory phenotype previously documented in vAT at this stage ^3,9^. Functional annotation localized many of these changes to oxylipin and eicosanoid signaling, glycerolipid signaling and storage, mitochondrial fatty-acid oxidation, membrane phospholipid metabolism and sphingolipid stress signaling (Figure S4C). Collectively, these results identify vAT as the principal site of late HFD-induced metabolic remodeling, characterized by coordinated disruption of lipid signaling, membrane-lipid composition and mitochondrial substrate handling.

### Endocannabinoid signaling is the leading enriched pathway in inflamed visceral fat

In scAT, metabolites and lipids decreased by 8 weeks of HFD feeding were enriched in pathways related to unsaturated fatty-acid biosynthesis, linoleic-acid metabolism, de novo fatty-acid biosynthesis and purine metabolism, indicating coordinated suppression of these metabolic processes relative to control-fed mice (Figure 2A; Table S2A). In vAT, the KEGG retrograde endocannabinoid-signaling pathway was the most significantly enriched pathway among metabolites and lipids increased after 8 weeks of HFD feeding (BH-adjusted P = 3 × 10⁻⁵; Figure 2B; Table S2B). The associated metabolic signature included 2-AG and multiple arachidonate-containing diacylglycerol and phosphatidylcholine species that may serve as substrates or precursors for endocannabinoid biosynthesis (Figure 2C; Table S1C)

**Figure 2.**
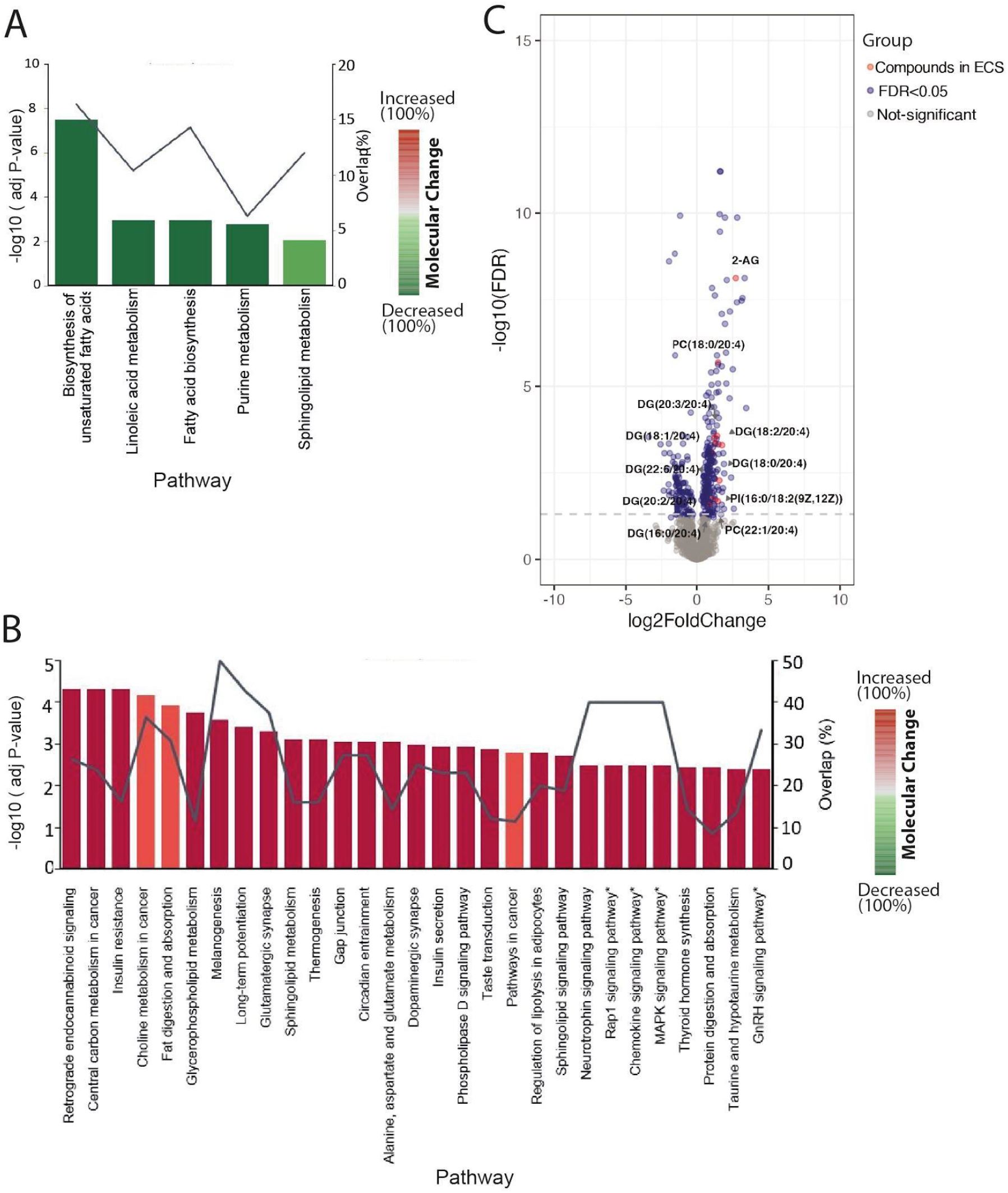
Effect of 8 weeks of HFD on the metabolome of visceral and subcutaneous adipose tissue. A-B) Pathway enrichment in scAT and vAT, respectively. Pathways required at least two mapped metabolites and BH-adjusted P < 0.05. The primary y axis shows -log10 (BH-adjusted P) and the secondary y axis shows pathway coverage. HFD selectively remodels the endocannabinoid system in visceral adipose tissue. C) Volcano plot of 2-AG and arachidonate-containing diacylglycerol and phosphatidylcholine species in vAT after 8 weeks of HFD.

Gene expression supported a coordinated visceral ECS response at week 8 (Figure S5 A-B; Table S3A-B). *Cnr2* and *Daglb* increased, whereas *Napepld, Faah*, and, more modestly, *Mgll* decreased. In scAT, *Daglb* and *Mgll* increased together, *Cnr1* decreased, and *Cnr2* and *Faah* remained unchanged. These opposing synthetic and degradative responses provide a plausible explanation for 2-AG accumulation in vAT (Table S3A) but not scAT (Table S3B). One week of HFD caused little ECS transcriptional remodeling apart from increased *Mgll* in vAT.

Promoter-associated chromatin marks changed in the same direction as several transcriptional responses. H3K27 acetylation and H3K4 monomethylation increased at two regions near the *Cnr2* promoter specifically in vAT at week 8. Increased *Daglb* expression was accompanied by enrichment of these marks at four promoter-proximal regions. The scAT reduction in *Cnr1* expression coincided with reduced H3K27 acetylation at its promoter (Figure S5 C-E; Table S3C). Together, the metabolite, transcript, and chromatin data indicate coordinated and depot-specific ECS remodeling.

### Targeted quantification confirms selective accumulation of 2 AG in visceral fat

Absolute quantification of several endocannabinoids by LC-MRM tandem mass spectrometry confirmed that HFD significantly increased 2-AG and Stearoylethanolamide (SEA) in vAT but not scAT (Figure 3A-B, S6, S7A-B). Whole-tissue AEA and Palmitoylethanolamide (PEA) did not change significantly in either depot (Figure 3A-D, S6, S7A-B).

**Figure 3.**
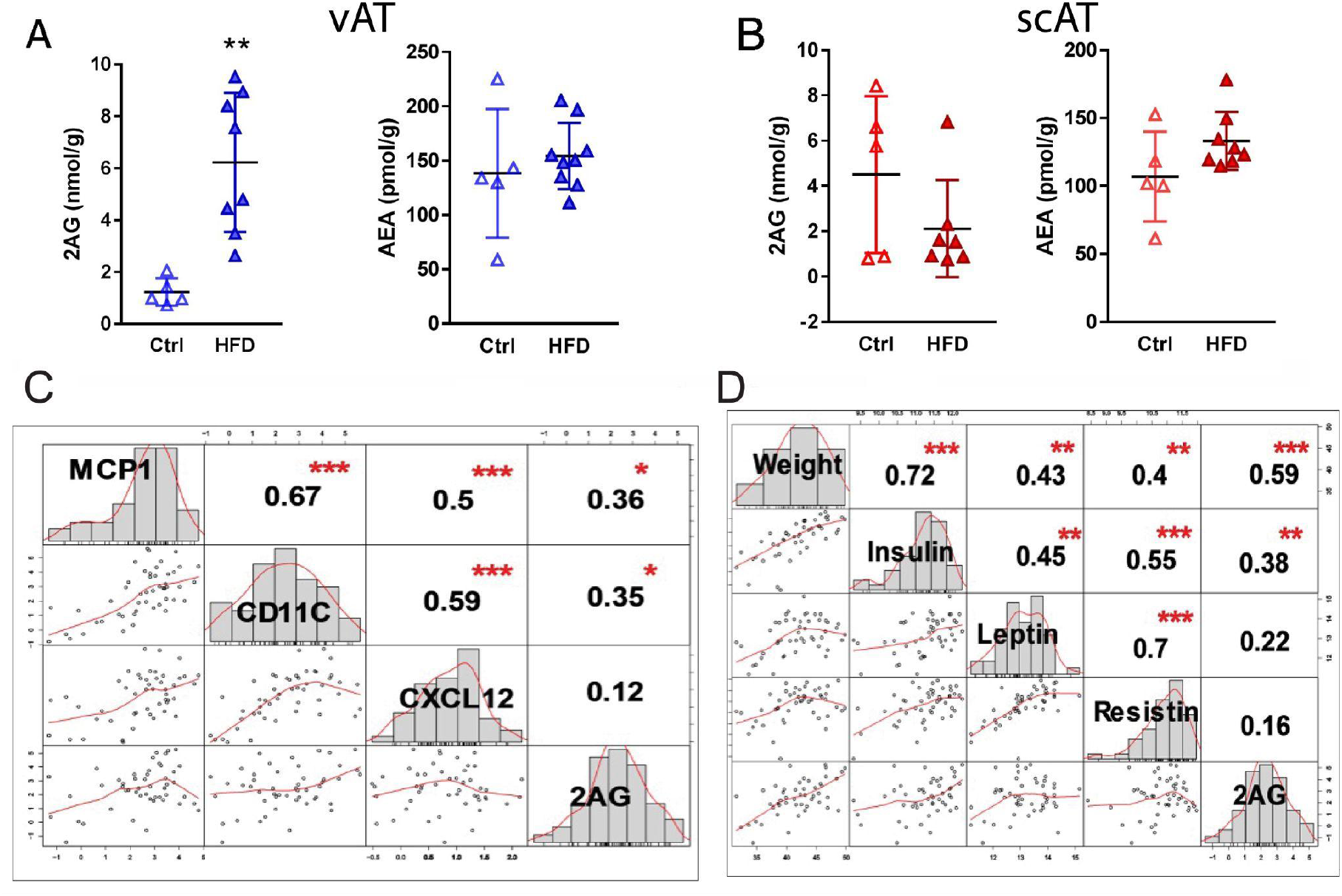
ECS quantification, its association with metabolic and inflammatory signals. A-B) Effect of HFD on the levels of the two main endocannabinoids: 2-AG and anandamide are measured in vAT (A) and scAT (B). Significant level in comparison with ctrl group has been notified as ** (p-value < 0.01). 2-AG levels in vAT correlated with inflammatory (C) and metabolic (D) signals. Significant correlations have been notified as *(p-value < 0.05), ** (p-value < 0.01), *** (p-value < 0.001), ****(p-value < 0.0001).

Visceral 2-AG concentrations correlated with the inflammatory and metabolic phenotype, including *Ccl2* and *Itgax* expression, circulating insulin, and body weight (Figure 3 C-D; Table S3D). Body weight showed the highest reported association (r = 0.52; Fisher z = 0.58; 95% CI 0.29-0.87).

Further indicating the intricate link between vAT inflammation and ECS, In low-responder HFD mice, induction of *Cnr2* and *Daglb*, and to a lesser extent *Dagla*, was absent, whereas downregulation of *Faah* and *Napepld* resembled that in good responders (Figure S7C-G). Thus, part of the ECS response tracked inflammatory status, while another component tracked diet exposure independently of notable inflammation.

### Adipocytes and the stromal vascular fraction contribute distinct components of the visceral ECS response

To define the cellular distribution of the ECS response, mature adipocytes and the stromal vascular fraction (SVF) were isolated from vAT and scAT after 8 weeks of control or HFD feeding (Figure 4A–H). In both depots, *Cnr1* expression was concentrated in adipocytes, whereas *Cnr2* was predominantly expressed in the SVF (Figure 4A–B). HFD selectively increased *Cnr2* expression in the vAT SVF, with no corresponding induction in scAT. Among genes involved in endocannabinoid metabolism, *Dagla*, *Napepld* and *Mgll* were expressed predominantly in adipocytes, whereas *Daglb* and *Faah* showed greater expression in the SVF; *Naaa* was more evenly distributed between fractions (Figure 4C–F). HFD altered the expression of 2-AG-metabolizing enzymes in a depot- and fraction-dependent manner. Notably, *Mgll*, which encodes the principal 2-AG-degrading enzyme, was induced in scAT adipocytes but not in vAT adipocytes, indicating a potential depot-specific difference in the capacity to limit 2-AG accumulation.

**Figure 4.**
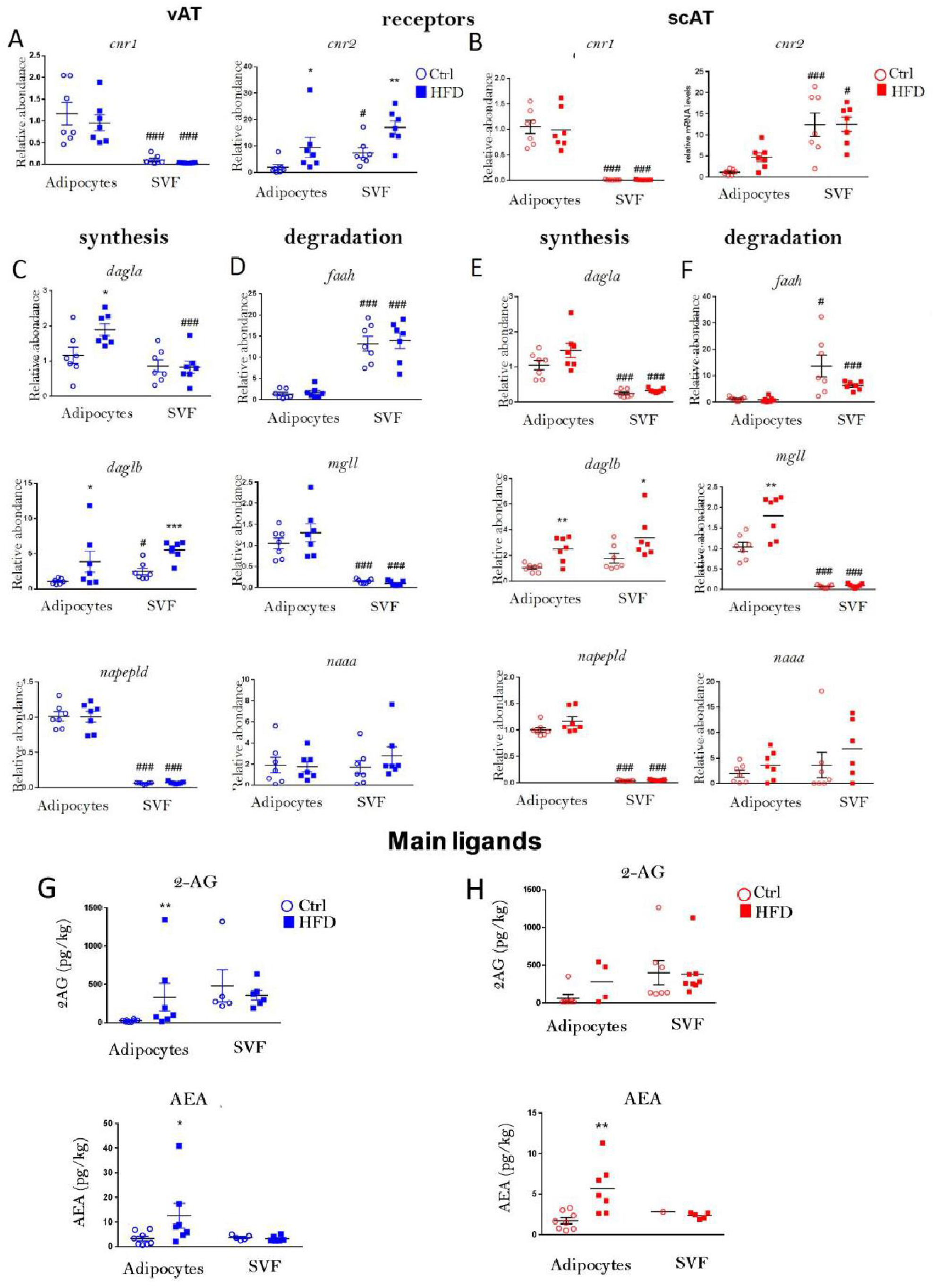
ECS regulation by HFD in visceral adipose tissue. mRNA levels of the indicated genes measured in separated adipocytes and SVF from vAT (A, C, D) and scAT (B, E, F) isolated from mice fed with HFD or chow diet for 8 weeks (n=8). 2-AG and AEA levels measured in separated adipocytes and SVF from vAT (G) and scAT (H) isolated from mice fed with HFD or chow diet for 8 weeks (n=4-5).* denotes the significant level of comparative analysis performed between HFD and chow diet as *(p-value < 0.05), ** (p-value < 0.01), *** (p-value < 0.001), ****(p-value < 0.0001) and ## notifying p-value < 0.001 showed the significance level of across fraction comparison for identical treatment.

Targeted lipid measurements supported this cellular distinction (Figure 4G–H). HFD increased 2-AG abundance in vAT adipocytes, whereas no corresponding accumulation was detected in scAT adipocytes or in the SVF of either depot. In contrast, AEA increased primarily in the adipocyte fraction of both vAT and scAT. These findings localize the excess visceral 2-AG pool to adipocytes while identifying the *Cnr2*-enriched vAT SVF as a distinct cannabinoid-responsive compartment.

The combined ligand and gene-expression profiles are consistent with compartmentalized ECS remodeling in which 2-AG accumulates in vAT adipocytes and may signal to *Cnr2*-expressing SVF cells, although direct paracrine signaling was not demonstrated.

### Cannabinoid selectively attenuates inflammatory signaling in activated adipocytes through macrophages and remodels their secretory profile

To investigate potential adipocyte–macrophage communication, RAW264.7 macrophages and differentiated 3T3-L1 adipocytes were examined both separately and in an insert-based co-culture system that permitted the exchange of soluble factors without direct cell contact (Figure 5A).

**Figure 5.**
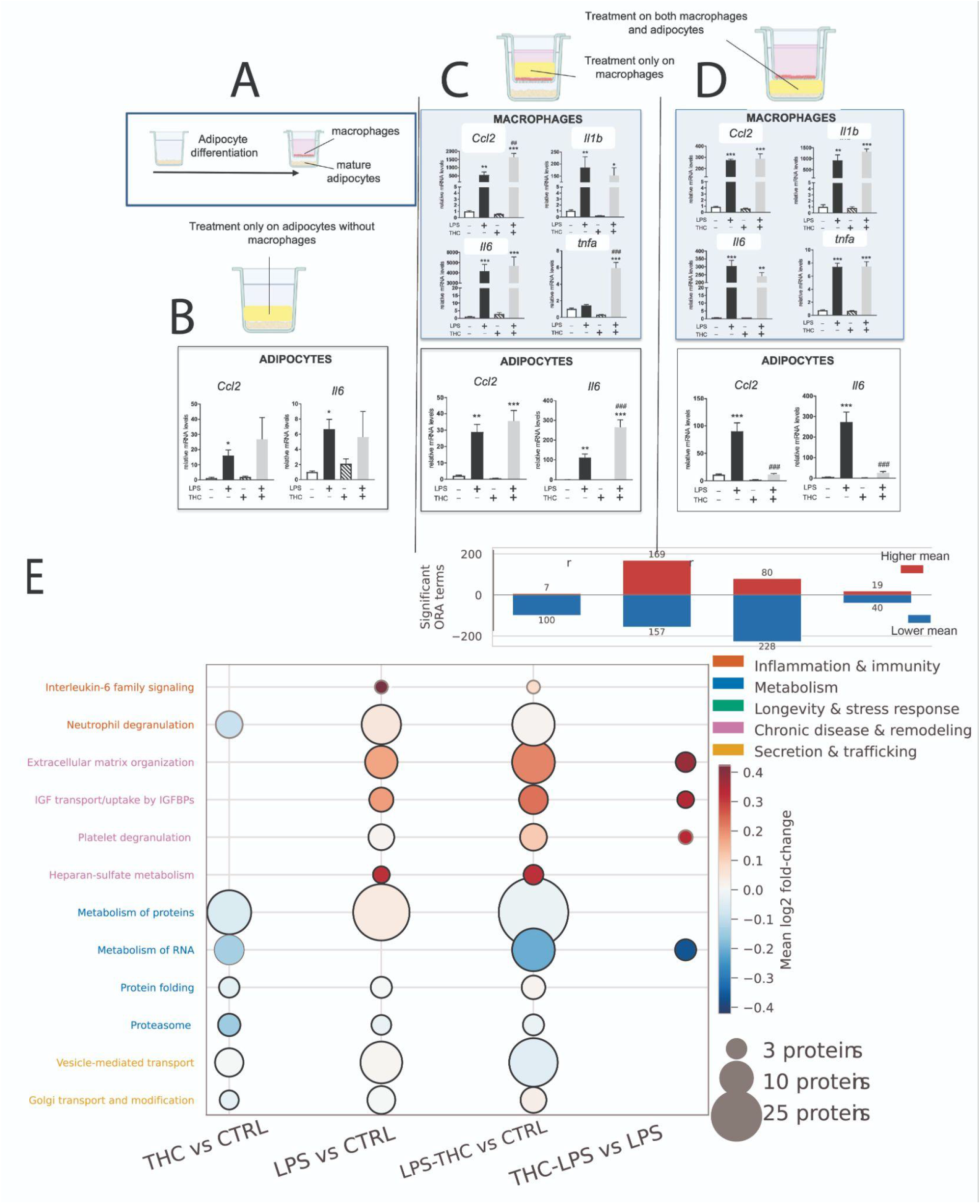
THC remodels the response of adipocytes and macrophages to LPS. (A) Experimental scheme to explore adipocyte macrophage cross-talk, RAW264.7 macrophages were plated in inserts and put in co-culture with differentiated 3T3-L1. (B) Differentiated 3T3-L1 adipocytes were treated with THC (5 µM) in combination with LPS (0.1 µg mL⁻¹) for 24 h. (C) RAW264.7 macrophages were put in inserts on top of mature adipocytes and a treatment with THC (5 µM) in combination with LPS (0.1 µg mL⁻¹) was performed only in macrophages for 24h. (D) RAW264.7 macrophages were put in inserts on top of mature adipocytes and both cell-types were treated with THC (5 µM) in combination with LPS (0.1 µg mL⁻¹) for 24h. mRNA levels of Tnfa, Il1b, Il6, and Ccl2 are shown as mean ± SE (n = 3). (E) Secreted protein profiling in adipocytes–macrophage co-cultures. Significantly enriched over-representation analysis (ORA) terms were highlighted according to the direction of their mean log₂ fold change. Representative enriched pathways were selected to summarize inflammatory signaling, extracellular-matrix remodeling, metabolism/proteostasis, and secretory trafficking while minimizing redundant pathway annotations.Only a subset of representative pathways is shown; the complete pathway enrichment profile is presented in Figure S9.

**Figure 6.**
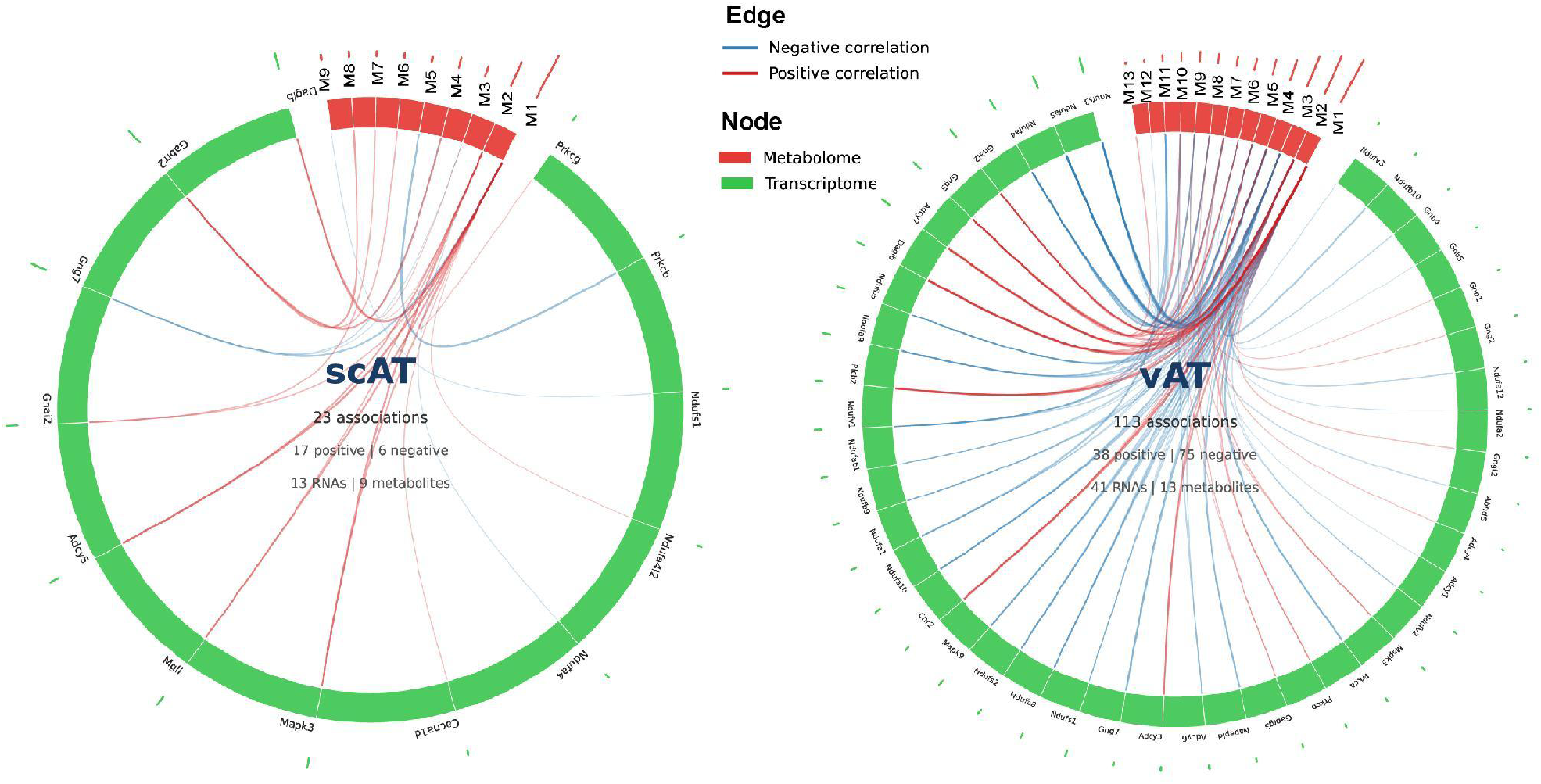
Functional activation of ECS pathway in vAT upon HFD. Cross-omics integration of endocannabinoid-system-associated transcripts and metabolites in adipose tissue. Transcriptomic and positive-ion-mode metabolomic features significantly altered by HFD in either scAT or vAT were mapped to the KEGG ECS signaling pathway and analyzed using sparse generalized canonical correlation analysis. Circos plots show transcript–metabolite associations with an absolute Sparse Generalized Canonical Correlation Analysis (sGCCA) correlation of |r| ≥ 0.90. Red edges indicate positive correlations and blue edges indicate negative correlations; edge thickness represents absolute correlation magnitude. The vAT network exhibited a predominance of strong negative transcript–metabolite associations, whereas the corresponding scAT associations were less extensive. The Cytoscape network displays only variables participating in at least one retained association and therefore represents a subset of the 118 transcripts and 13 metabolites entered into the integration analysis.

Because 2-AG is rapidly degraded under culture conditions, delta-9-tetrahydrocannabinol (THC) was used as a more stable pharmacological activator of cannabinoid signaling. THC had little effect on basal expression of *Tnfa*, *Il1b*, *Il6* or *Ccl2* in either cell type (Figure S8) and did not influence adipocyte response to pro-inflammatory LPS stimulation (Figure 5B). In contrast, in co-culture experiments, THC dampened the pro-inflammatory effect of LPS in adipocytes, but only in presence of macrophages, suggesting that the adipocyte-macrophage cross-talk is required to mediate the anti-inflammatory effect of THC through the secretion of extracellular mediators (Figure 5C-D).

Secretome profiling provided an additional measure of this cross-talk (Figure 5E, S9). Because the conditioned medium was collected from a co-culture containing both cell types, the detected proteins represent a composite adipocyte–macrophage secretome and cannot be assigned to either cellular source. Rather than broadly reversing the LPS-induced secretory program, THC produced a selective redistribution of secreted-protein pathways (Table S4A-B). In the direct THC-LPS versus LPS comparison, proteins associated with extracellular-matrix organization, ECM proteoglycans, integrin interactions, IGF transport and platelet-associated secretion showed higher mean abundance profiles. Conversely, proteins assigned to RNA metabolism and related protein-processing pathways showed lower profiles (Figure S9; Table S4B). Together with the cell-specific transcriptional measurements, these findings suggest that THC modifies the inflammatory cross-talk between macrophages and adipocytes while reorganizing the extracellular and secretory environment of the co-culture. However, the present analysis does not establish which cell type produced the altered proteins or whether the resulting secretome promotes inflammatory resolution, tissue repair, fibrosis or another remodeling outcome.

### Integrated ECS remodeling is associated with reduced mitochondrial gene expression in visceral fat

Integration of ECS-related metabolites with RNA-seq data revealed denser metabolite-gene correlations in vAT than in scAT. Several genes encoding mitochondrial respiratory-chain subunits, including members of the *Nduf* family, were negatively associated with ECS activation in vAT (Table S5). Previous work shows that CB1 stimulation can suppress AMPK signaling and regulators of mitochondrial biogenesis in white adipocytes ^12–14^. The present data therefore raise the possibility that visceral ECS activation contributes to reduced adipocyte mitochondrial activity, although the correlations do not establish directionality or cell type.

Strong positive correlation has been observed between chemical compound of ECS and several genes such as *Adcy5, Cacna1d*, *Daglb, Gabrr2, Gnai2, Mapk3, Mgll, Mapk3, Prkcg and Ndufa4l2*. In contrast, *Ndufs1, Ndufa4, Gng7,* and *Prkcb* are shown to be negatively correlated with metabolites contributing to ECS.

In scat; M1= DG(18:0/20:4/0:0), M2= L.Glutamic.acid, M3= 2.Arachidonylglycerol (2-AG), M4= DG(18:0/20:4/0:0), M5= Cyclic.AMP, M6= DG(18:2/20:4/0:0), M7= DG(18:0/20:4/0:0)-2, M8= DG(20:4/22:6/0:0), M9= DG(20:3n6/0:0/20:4n6). In vAT; M1= DG(18:0/20:4/0:0), M2= DG(20:3n6/0:0/20:4n6), M3= PC(18:0/20:4), M4= DG(20:4/22:6/0:0), M5= DG(18:0/20:4/0:0)-2, M6= DG(16:0/20:4/0:0), M7= DG(18:0/20:4/0:0), M8= PC(22:4/18:0), M9= L.Glutamic.acid, M10= DG(18:2/20:4/0:0), M11= Cyclic.AMP, M12= PC(22:1/20:4), M13= 2.Arachidonylglycerol (2-AG). ECS= Endocannabinoid system.

## Discussion

This study identifies the ECS as a prominent component of the metabolic remodeling that accompanies the emergence of inflammation in vAT during sustained HFD feeding. At 8 weeks, ECS signaling was among the highest-ranked enriched metabolic pathways in vAT and was supported by concordant changes across metabolomic, transcriptomic, promoter-associated chromatin and cellular-fraction measurements. The response was not a generalized increase in endocannabinoids across adipose depots. Instead, it was characterized by selective accumulation of 2-AG in vAT, extensive remodeling of ECS-related genes and a much denser transcript–metabolite association network in vAT than in scAT. These findings position the ECS within a broader, depot-specific adaptation to prolonged nutrient excess.

Metabolic changes were detectable in vAT before the development of the pronounced inflammatory phenotype observed at 8 weeks. After only 1 week of HFD feeding, relatively few metabolites were significantly altered, but vAT already showed changes associated with sphingomyelin metabolism. These findings may represent an early molecular response during a period in which conventional inflammatory markers remain minimally altered. However, describing this response as an “ultra-early warning system” would require prospective evidence that the metabolites predict subsequent inflammation at the individual-animal level. The present data instead support the more limited conclusion that lipid remodeling precedes obvious vAT inflammation and may participate in, or reflect, the early adaptation to HFD. The marked expansion of the metabolic signature by week 8 indicates that most ECS-associated remodeling develops during sustained dietary exposure rather than immediately after diet initiation.

The selective increase in vAT 2-AG is consistent with clinical evidence linking peripheral ECS activity to visceral adiposity. In humans, circulating 2-AG is higher in visceral than in subcutaneous obesity and correlates with visceral fat mass, fasting insulin and reduced insulin sensitivity ^8^. Moreover, lifestyle-induced reductions in visceral adiposity are accompanied by substantial decreases in plasma 2-AG, and these changes correlate with improvements in triglyceride-related metabolic risk^15^. Our results extend these observations by identifying visceral adipocytes as the principal cellular compartment in which excess 2-AG accumulates after HFD feeding. The coordinated changes in 2-AG-metabolizing enzymes and the selective accumulation of 2-AG in vAT adipocytes identify visceral adipose tissue as a plausible contributor to elevated circulating 2-AG in obesity. However, because tissue-to-plasma 2-AG flux was not directly measured, the relative contribution of vWAT compared with other organs remains unresolved. Nevertheless, the expansion of visceral fat mass together with increased adipocyte 2-AG provides a plausible basis for investigating whether vAT contributes to elevated circulating endocannabinoids in obesity.

The divergence between vAT and scAT further demonstrates why the abundance of a single synthetic or degradative enzyme is insufficient to predict tissue endocannabinoid concentrations. HFD altered genes involved in 2-AG and AEA metabolism, including *Dagla, Daglb, Mgll, Abhd6*, *Abhd12, Napepld* and *Faah*, but the direction and cellular distribution of these changes differed between depots. Both tissues showed induction of genes supporting 2-AG synthesis, whereas scAT adipocytes additionally induced *Mgll,* potentially increasing their capacity for 2-AG hydrolysis. This matched degradative response was absent from vAT adipocytes, in which 2-AG accumulated. Endocannabinoid concentrations therefore appear to reflect the combined effects of synthetic and degradative capacity, substrate availability, enzyme activity and cell-specific lipid flux. This interpretation is consistent with evidence that insulin resistance disrupts the coordinated regulation of ECS-metabolizing enzymes in adipocytes and is associated with increased adipose and circulating endocannabinoids ^16^.

The cellular-fraction data reveal an additional level of organization. Under control conditions, *Cnr1* was mainly expressed in adipocytes, whereas *Cnr2* was enriched in the SVF. HFD selectively increases *Cnr2* in the vAT SVF, the compartment containing immune, endothelial, progenitor and other stromal cells. In parallel, excess 2-AG localized to the vAT adipocyte fraction. This complementary distribution is consistent with a model in which adipocyte-derived lipid mediators signal to cannabinoid-responsive SVF cells. However, increased *Cnr2* cannot yet be assigned to a particular immune-cell population, because the SVF is heterogeneous and changes in its expression profile may reflect both transcriptional regulation and HFD-induced shifts in cellular composition. Flow cytometry, spatial analysis or single-cell profiling will be necessary to identify the relevant *Cnr2*-expressing cells and establish whether adipocyte-derived 2-AG reaches them at biologically active concentrations.

The cell experiments provide functional evidence that cannabinoid stimulation can modify an established inflammatory response, but they do not identify the responsible mediator. In presence of both cells, adipocytes and macrophages, THC reduced LPS-induced *Il6* and *Ccl2* expression in differentiated adipocytes without significantly influencing macrophage inflammatory response. The cross-talk between these two cell types mediated therefore a selective effect on adipocyte inflammatory outputs, rather than triggering a generalized immunosuppression. Reduction of *Ccl2* could be relevant to signals governing monocyte recruitment ^17^, while attenuation of *Il1b* could limit amplification of local inflammatory signaling ^18^. Nevertheless, THC activates both CB1 and CB2 and may engage additional targets; it is also not pharmacologically equivalent to endogenous 2-AG. The results consequently cannot be assigned specifically to CB2 or used alone to conclude that adipocyte-derived 2-AG suppresses macrophage inflammation.

The present data are most consistent with cannabinoid-dependent attenuation of selected responses in already activated adipocytes, rather than with a universal anti-inflammatory function of CB2.

Secretome profiling broadened this interpretation beyond inflammatory transcripts. THC did not globally reverse the LPS-induced secretory response. Instead, addition of THC to LPS-stimulated co-culture selectively altered the relative abundance of secreted proteins assigned to inflammatory, extracellular-matrix and intracellular-signaling pathways. Proteins associated with ECM organization, proteoglycans, integrin interactions, IGF transport and regulated secretion showed higher mean-abundance profile, whereas proteins related to RNA metabolism and protein processing showed lower profiles. The lower extracellular abundance of proteins annotated to RNA metabolism and protein processing should not be interpreted as direct suppression of these intracellular pathways. Because the analyzed conditioned medium was collected from the adipocyte–macrophage co-culture, the cellular origin of individual proteins cannot be assigned from the bulk proteomic data. The extracellular proteins may include conventionally secreted factors as well as proteins released through extracellular vesicles, ectodomain shedding or unconventional secretion ^19^ ^20^. Indeed, intracellular proteins involved in RNA processing, transcription and translation can be detected extracellularly as a consequence of cellular leakage^21^. The observed reduction could therefore reflect altered release or extracellular-vesicle transport, reduced cell damage, or differences in cellular abundance rather than reduced intracellular RNA-processing activity. Distinguishing among these possibilities will require parallel measurements of cell viability, intracellular protein abundance and extracellular-vesicle-associated proteins.

The cross-omics analysis further emphasized the greater scale of ECS-associated remodeling in vAT. Negative associations predominated in vAT, whereas the smaller scAT network was primarily positive. Among the negatively associated transcripts were multiple *Nduf* genes encoding subunits or components associated with mitochondrial complex I ^22^. This pattern links increased ECS-related lipid features to reduced expression of genes supporting oxidative phosphorylation, but it does not demonstrate inhibition of mitochondrial respiration. Direct measurements of oxygen consumption, complex I activity, mitochondrial abundance and reactive oxygen species are nevertheless required before the vAT network can be interpreted as functional mitochondrial impairment.

Taken together, the results support a compartmentalized model of ECS remodeling in inflamed visceral adipose tissue. In this model, sustained HFD exposure increases 2-AG availability in adipocytes while expanding a *Cnr2*-enriched SVF compartment. Cannabinoid signaling could consequently have different effects in neighboring cell populations: CB1-associated signaling in adipocytes may contribute to lipid storage and reduced oxidative capacity, whereas cannabinoid-responsive immune cells may selectively attenuate some inflammatory outputs while modifying extracellular and secretory programs. These actions could coexist, potentially explaining why systemic ECS overactivity is associated with obesity even though cannabinoid stimulation can suppress selected macrophage responses. This dual-action model remains a testable hypothesis rather than a demonstrated causal mechanism.

Several limitations define the scope of these conclusions. First, the temporal ECS pattern and, at 8 weeks, the difference between low- and high-inflammation HFD-fed mice establish an association of ECS with inflammatory status but do not show that ECS remodeling initiates inflammation. Second, transcript abundance and promoter-associated chromatin marks indicate regulatory capacity, not enzyme activity or ligand flux. Third, adipocyte/SVF fractionation is vulnerable to variable recovery and cross-contamination, and the heterogeneous SVF prevents assignment of Cnr2 induction to a defined cell type. Fourth, THC is not receptor selective, and the macrophage and adipocyte cell lines do not reproduce the full cellular environment of primary adipose tissue. Fifth, secretome pathway enrichment does not establish the biological activity of the released proteins. Finally, the Sparse Generalized Canonical Correlation Analysis (sGCCA) networks represent strong correlations rather than direct biochemical interactions or causal regulatory relationships.

These findings could subsequently be validated using cell-type-specific *Cnr2* deletion in vivo. Stable-isotope tracing could determine ligand production, turnover and release, while spatial or single-cell analyses could resolve the identities and states of cannabinoid-responsive SVF cells. Direct analysis of adipocyte mitochondrial respiration and ROS production would establish whether the inverse association between ECS metabolites and complex I transcripts has functional consequences. These experiments may also inform therapeutic strategies that modulate pathological ECS tone without disrupting central cannabinoid signaling, for example, peripherally restricted CB1 blockade or local modulation of endocannabinoid synthesis and degradation. Such approaches may offer greater precision than global receptor blockade, but their value will depend on preserving potentially adaptive immune effects while limiting the metabolic consequences of excessive adipocyte ECS activity.

## Materials and Methods

### Animals and dietary intervention

All experiments involving animals were approved by the Veterinary Office of the Canton Vaud (Switzerland) in accordance with the Federal Swiss Veterinary Office Guidelines and conform to the Commission Directive 2010/63/EU (VD-2942.b). Male C57BL/6 mice were purchased from Janvier Labs and housed five per cage. Four-week-old mice received a 10% fat control diet (D12450J, Research Diets) for 2 weeks. At 6 weeks of age, mice were either switched to a high-fat diet (HFD) containing 60% fat (D12492, Research Diets) or maintained on the control diet for 1 or 8 weeks. Random blocking was used for experimental allocation. The progression of diet-induced obesity was monitored by regular measurements of body weight (Figure S1A). The metabolomics cohort included 116 mice. Mice were maintained on a 12-hour light and 12-hour dark cycle with food and water ad libitum and were euthanized with carbon dioxide between zeitgeber time 2 and 5. Perigonadal fat was analyzed as vAT and inguinal fat as scAT. Tissues were weighed, aliquoted, and stored at -80 degrees Celsius.

### Phenotypic definition of inflammatory response

Body weight and circulating insulin, leptin, and resistin were combined with vAT expression of *Ccl2*, Itgax, and *Cxcl12*. Principal-component analysis and partitioning around medoids were used to characterize variation within diet groups. Macrophage infiltration was assessed by F4/80 staining. HFD-fed animals with a phenotype close to control animals and limited vAT macrophage infiltration were classified as low inflammation (Low-INFL), as opposed to the other HFD-fed animals developing the expected vAT inflammation ^3^.

### Untargeted metabolomics Sample preparation

A total of 264 analytical injections, comprising 232 adipose-tissue extracts, represented visceral adipose tissue (vAT) and subcutaneous adipose tissue (scAT) collected from 116 mice and 32 pooled quality-control samples using established protocol ^10,23,24^ adipose-tissue samples were processed in randomized order. For each sample, 10–20 mg of frozen visceral or subcutaneous adipose tissue was extracted with 400 µL of ice-cold methanol/ethanol/water (2:2:1, v/v/v).

Blank extractions were processed with the samples. A pooled biological quality-control sample was prepared by combining 4 mg of tissue from each biological sample and was analyzed repeatedly throughout the analytical sequence.

### Experimentation

Untargeted metabolomics was performed using an established platform ^10,23,24^. Briefly, the UPLC (Dionex,Thermo Fisher Scientific) hyphenated with HRMS QExactive plus (Thermo ScientificTM Q ExactiveTM) has been used. Metabolome profile of fat tissue (i.e. v-AT and sc-AT) was obtained for 264 samples including 32 QCs and 232 adipose tissue. HRMS was interfaced with an electrospray ionization (ESI) source operated in both negative (NEG) and positive (POS) polarities. UPLC was performed on a C18 Kinetex, 2.6 μm, 50 mm × 2.1 mm I.D. column (Phenomenex, PA) in Reverse Phase chromatography ^10,23,24^. Raw data were then transformed to mzXML format using MSConvert (Proteo Wizard 3.07155) and pre-processed for peak peaking, chromatogram alignment and isotope annotation using open access XCMS online (https://xcmsonline.scripps.edu). XCMS runs on UPLC-QExactive parameters by setting peak detection on Centwave. Preprocessing generated 21,055 features across positive and negative modes; filtering and annotation retained 2,030 features. Analytical drift and batch effects were evaluated by principal-component analysis, hierarchical clustering, and feature-wise adjusted coefficients of determination, then corrected using the ber model implemented in DBnorm ^10,23,24^. Protein normalization has been used for signal correction.

### Targeted metabolomics

#### Sample preparation and endocannabinoid quantification

AEA, 2-AG, PEA, OEA, and SEA were quantified using their corresponding isotope-labeled internal standards: AEA-d4, 2-AG-d5, PEA-d3, OEA-d2, and SEA-d3. The extraction solvent consisted of methanol/ethanol/water (2:2:1, v/v/v).

For intact adipose tissue, approximately 30 mg of sample was extracted with 500 µL of extraction solvent containing 20 ng mL⁻¹ each of AEA-d4, PEA-d3, OEA-d2, and SEA-d3 and 200 ng mL⁻¹ 2-AG-d5. Samples were homogenized with three stainless-steel beads for 5 min and centrifuged at 14,700 × g for 10 min at 4 °C. A 450-µL aliquot of supernatant was transferred to a clean tube and evaporated to dryness. The residue was reconstituted in 100 µL of methanol/water (70:30, v/v), producing final internal-standard concentrations of approximately 90 ng mL⁻¹ for AEA-d4, PEA-d3, OEA-d2, and SEA-d3 and 900 ng mL⁻¹ for 2-AG-d5, assuming complete recovery. The residual tissue pellet was retained for protein measurement.

Because isolated adipocytes were collected in larger-volume tubes, each adipocyte preparation was extracted with 1.5 mL of solvent containing 6 ng mL⁻¹ each of AEA-d4, PEA-d3, OEA-d2, and SEA-d3 and 60 ng mL⁻¹ 2-AG-d5. Following homogenization and centrifugation, 1 mL of supernatant was evaporated to dryness and reconstituted in 100 µL of methanol/water (70:30). The resulting internal-standard concentrations were approximately 60 ng mL⁻¹ for AEA-d4, PEA-d3, OEA-d2, and SEA-d3 and 600 ng mL⁻¹ for 2-AG-d5.

Calibration standards containing AEA, 2-AG, PEA, OEA, and SEA were prepared in methanol and spiked with fixed amounts of their corresponding isotope-labeled internal standards. The standards were evaporated to dryness and reconstituted in 50 µL of methanol/water (70:30, v/v). Internal-standard concentrations in the calibration samples were matched to those in the corresponding tissue or adipocyte extracts.

Preparation of calibration curve:

A- Standard (AEA, 2-AG, PEA, OEA, SEA) at 1 µg/ml. 100 µL of each in 1mL MeOH

B- Standard of 5-IS at 100 ng/ml. Dilute A 10 times.

C- Standard 5-IS at 10 ng/ml. Dilute B 10 times.

D- Standard 5-IS at 1 ng/ml. Dilute C 10 times.

|  | A |  |  | B |  |  | C |  |  | D |  |  |
| --- | --- | --- | --- | --- | --- | --- | --- | --- | --- | --- | --- | --- |
| Volume(µL) | 25 | 50 | 150 | 100 | 200 | 400 | 100 | 200 | 500 | 10 | 50 | 100 |
| Conc( ng/mL) | 500 | 1000 | 3000 | 200 | 400 | 800 | 20 | 40 | 100 | 0.2 | 1 | 10 |

### Experimentation

To measure AEA, 2-AG, PEA, OEA, and SEA, dried residue was reconstituted in methanol and water at 7:3. Quantification used an established LC-MRM/MS method on a QTRAP 5500 hybrid triple-quadrupole linear ion-trap instrument coupled to a Dionex Ultimate 3000 LC system in positive electrospray mode^25–27^. Separation used a Chromolith RP18 column with an acetonitrile-water gradient containing 0.1% formic acid at 0.4 mL per minute. Eleven-point calibration ranges were 0.1-5 ng per mL for AEA, palmitoylethanolamide, and oleoylethanolamide and 1-30 ng per mL for 2-AG.

Analytical carry-over was below 0.01%, and the chromatographic method resolved the target endocannabinoids within a 3- to 5-minute elution window.

### Adipocyte and stromal vascular fraction isolation

For fractionation, 32 mice were assigned to control diet or HFD for 8 weeks, with eight animals per depot and diet group. Fresh vAT and scAT were digested for 30 minutes at 37 degrees Celsius in HEPES-buffered saline containing 1.5% bovine serum albumin, 5 mM glucose, 1.3 mM calcium chloride, and 0.2% collagenase II. Material was filtered through a 100-micrometer mesh and incubated on ice. The infranatant was passed through a 30-micrometer mesh and centrifuged at 700 x g for 10 minutes at 4 degrees Celsius to collect SVF. Floating mature adipocytes were washed by repeated settling and removal of infranatant. Both fractions were snap frozen.

### Cell culture and inflammatory stimulation

3T3-L1 preadipocytes (American Type Culture Collection, Manassas, VA) were cultured in DMEM (Gibco #31966-021, Thermofisher) supplemented with 10% bovine calf serum (Gibco #26010-074, Thermofisher). Two days after confluence, differentiation was induced with fetal calf serum (Biowest #S1810-500, Nuaillé, France), 1µM dexamethasone (Sigma-Aldrich #D2915), 0.5mM 3-isobutyl-1-methylxanthine (IBMX -Sigma-Aldrich, #I7018), and 10µg/ml insulin (Sigma-Aldrich #I2643). The medium was replenished every other day by keeping only insulin.

RAW264.7 macrophages (American Type Culture Collection, Manassas, VA) were cultured in DMEM with 10% fetal calf serum. For co-culture experiments 3T3-L1 adipocytes were differentiated for 9 days in Falcon Companion Plates, then macrophages were plated in inserts (Falcon 353095) and cells were treated with Lipopolysaccharide (LPS – Sigma Aldrich, #L4391, dissolved in Phosphate buffer saline) and/or tetrahydrocannabinol (THC Lipomed #THC-135-1LE, dissolved in methanol) at the indicated concentrations in DMEM without serum. After 24h cell medium and cells were frozen at -80°C for secretome and gene expression analysis, respectively.

### Secretome proteomics

Conditioned medium was pooled from three wells per sample and concentrated with 30-kDa Amicon Ultra-4 filters. Proteins were reduced with tris(2-carboxyethyl)phosphine, alkylated with iodoacetamide, digested overnight with trypsin, cleaned on C18 material, and supplemented with indexed retention-time peptides. Peptides were analyzed on an Orbitrap Fusion Lumos coupled to an Easy-nLC 1200 system. Data-dependent acquisitions generated a mouse secretome spectral library, and data-independent acquisitions were searched and quantified with Spectronaut version 14.8. The library contained 6,151 proteins at 1% protein FDR; the DIA analysis quantified 3,460 proteins at 1% peptide and protein FDR. Treatment effects were assessed by ANOVA followed by Tukey HSD correction.

### Gene expression and multiomics integration

For the animal experiment with adipocyte-SVF separation and for cell experiments, RNA was purified with RNeasy columns and reverse transcribed with ImProm-II. Quantitative PCR used Kapa Probe Fast chemistry on an Applied Biosystems real-time PCR system. Technical triplicates were averaged after reproducibility checks. Relative expression was calculated by the 2^-delta-delta-Ct method using Tbp for tissue and Hprt for cell experiments, as housekeeping genes. RNA- and ChIP-seq data of the main animal experiment were retrieved from GEO, accession number GSE132885. Metabolite, RNA-seq, and histone-mark datasets were mapped to KEGG ECS signaling components for integrative correlation analysis. H3K27 acetylation was used as a mark of active enhancers and H3K4 monomethylation as a broader enhancer-associated mark.

### Statistical and Pathway analysis

Two-tailed Student t tests were used for pairwise HFD-versus-control gene-expression comparisons. Significance thresholds were reported as P below 0.05, 0.01, 0.001, or 0.0001. Pearson correlation coefficients quantified associations between vAT 2-AG and metabolic or inflammatory measures; 95% confidence intervals were obtained after Fisher z transformation. Untargeted metabolite and pathway thresholds are described above. Sample sizes were eight animals per group for fraction gene-expression analyses, four to five per group for fraction endocannabinoid measurements, and three independent replicates for cell-culture experiments.

LIMMA was used for depot- and time-specific differential analysis on Log2 transformed data. Putative identities were assigned by accurate-mass matching to the Human Metabolome Database, evaluated against retention behavior and peak shape, and confirmed by MS2 spectra (Table S1). Differential metabolites required an absolute log2 fold change above 1 and Benjamini-Hochberg adjusted P below 0.05.

Pathway over-representation analysis was performed against the ConsensusPathDB database (http://cpdb.molgen.mpg.de) using a hypergeometric test with multiple-testing correction. Pathways were considered significant if they contained at least two mapped metabolites and had a q-value <0.05.

## Data and Code Availability

RNA- and ChIP-seq data of the main animal experiment were retrieved from our previous study^3^ (Caputo et al 2021, GEO accession number GSE132885). The DBnorm R package^10^ used to visualize and correct analytical drift is available at https://github.com/NBDZ/dbnorm, and its methodological evaluation is available as a bioRxiv preprint [9].

## Ethics Statement

All mouse procedures were approved by the Swiss Veterinary Office under authorization numbers VD-2942.b and VD-3378.

## Author Contributions

N.B. performed the untargeted and targeted metabolomics analyses and integrated these results with collaborator-generated phenotypic, transcriptomics, and epigenomic data. Contributed in tissue and cell experiments. T.C. performed animal experiments and generated RNA and epigenomic data, T.S. performed secretome analysis, C.W. contributed to the in vivo experiments. N.G. and B.D. participated in the initial elaboration of the project and data discussion. F.G. and A.T. conceived and supervised the study. N.B., F.G. and A.T. wrote the manuscript.

## Competing Interests

No competing interests were stated in the source thesis.

## Acknowledgements

This research was supported by Swiss National Science Foundation [grants #310030-156771 and #31003A-182420]. Animal work was performed at the Center for Integrative Genomics at the University of Lausanne.

## Supplemental Table Legend

**Table S1. Metabolome analysis of AT**

**Table S1A- Metabolite analysis, cross diet treatment, time point and tissue depots**

**Table S1B. Metabolite classified captures across diet treatment and time point across in scAT**

**Table S1C. Metabolite classified captures across diet treatment and time point across in vAT**

**Table S2. Differential results of vAT metabolome analysis**

**Table S2A:** List of metabolites showing significant changes associated with 8 weeks of HFD treatment in scAT. Significance threshold defined as log2FC >1 (FC>2 or FC<0.5) and BH adj-p-value <0.05. List of significant metabolites was determined in both positive and negative polarities and Identifications status determined if a metabolic feature confirmed by MSMS spectra or by peak shape & retention time denoted as “semi”. Retention time (RT) detriments the time a metabolite pass through coulumn bases on our RPUPLC condition. Chemical class of compound is also presented.

The level of significance for these metabolites have been also presented for other compared groups, 8week-HFD vs Ctrl in vAT, 1week-HFD vs Ctrl in scAT, 1week-HFD vs Ctrl in vAT, 8week Ctrls vs Ctrl in vAT, 8week Ctrls vs Ctrl in scAT, 8-week vAT vs scAT in low responders (LR), 8-week High responders (HR) vs low responders (LR) in vAT and also scAT, 8-week LR vs Ctrl in vAT and scAT, 1-week vAT compared to scAT in Ctrls and 8-week vAT compared to scAT in Ctrls.

**Table S2B:** List of metabolites showing significant changes associated with 8 weeks of HFD treatment in vT. Significance threshold defined as log2FC >1 (FC>2 or FC<0.5) and BH adj-p-value <0.05. List of significant metabolites was determined in both positive and negative polarities and Identifications status determined if a metabolic feature was confirmed by MSMS spectra or by peak shape & retention time denoted as “semi”. Retention time (RT) detriments the time a metabolite pass through a column based on our RPUPLC condition. A chemical class of compounds is also presented.

The level of significance for these metabolites have been also presented for other compared groups, 8week-HFD vs Ctrl in vAT, 1week-HFD vs Ctrl in scAT, 1week-HFD vs Ctrl in vAT, 8week Ctrls vs Ctrl in vAT, 8week Ctrls vs Ctrl in scAT, 8-week vAT vs scAT in low responders (LR), 8-week High responders (HR) vs low responders (LR) in vAT and also scAT, 8-week LR vs Ctrl in vAT and scAT, 1-week vAT compared to scAT in Ctrls and 8-week vAT compared to scAT in Ctrls.

**Table S3. Genes and associated histone markers of the endocannabinoid system (ECS) and correlations between 2-AG levels and metabolic and inflammatory markers of obesity.**

Table S3A. Gene expression of ECS in vAT Table S3B. Gene expression of ECS in scAT

Table S3C. Histone marks at promoter region of ECS genes

Table S3D. Correlation coefficient and significance level between 2-AG and markers of inflammation (C) and clinical markers of obesity (D) in vAT; data contains information from 5 mice from control group, 33 in the HFD group. Control and HFD groups chosen from 8 weeks feeding period.

Note1: Log2 transformed data for inflammatory markers and plasmatic level of Insulin, Leptin and Resistin are considered.

**Table S4. Secretome analysis in Adipocyte-machrophages coculture.**

Table S4A- Differential analysis of secreted proteins in Adipocyte-macrophage coculture

Table S4B- Pathway ORA analysis of proteins differentially expressed at each treatment.

**Table S5- Correlation between RNAseq and metabolome of ECS Supplemental Figure Legend**

## Supplemental Figure Legend

**Figure S1.**
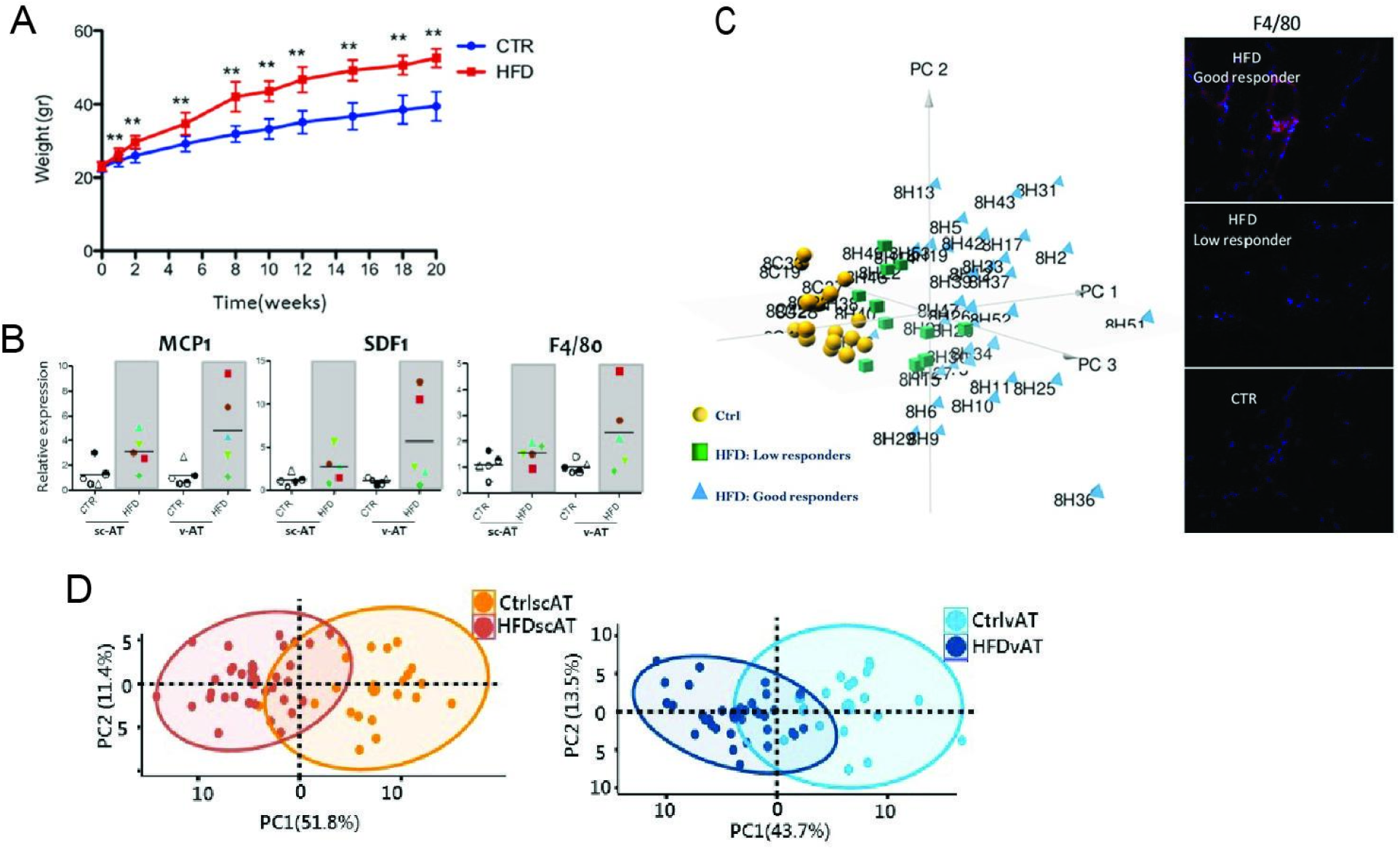
Efficacy of HFD in adipose tissue. HFD induced (A) obesity and (B) inflammation in visceral, but not in subcutaneous adipose tissue. C) Clustering of mice based on the features of phenotype. Right panel) PCA analysis demonstrated by features of obesity and inflammation in the mice that received either HFD or Ctrl Diet for 8 weeks. Clusters are represented by different colors. Mice receiving Ctrl Diet are presented by yellow circle points, mice included in HFD-fed are represented by blue triangles and the HFD-fed mice which showed a Low-Inflammation (Low-INFL) phenotype are represented by green rectangles. Right panel) vAT F4/80 staining of representative subset of three clusters. Low responders correspond to Low-INFL mice that developed low inflammatory response upon HFD treatment, while good responders developed significant induction of inflammation upon HFD. D) PCA analysis of raw data. Differential metabolite and explained variation on scAT (left) and vAT (right) metabolome on week 8.

**Figure S2.**
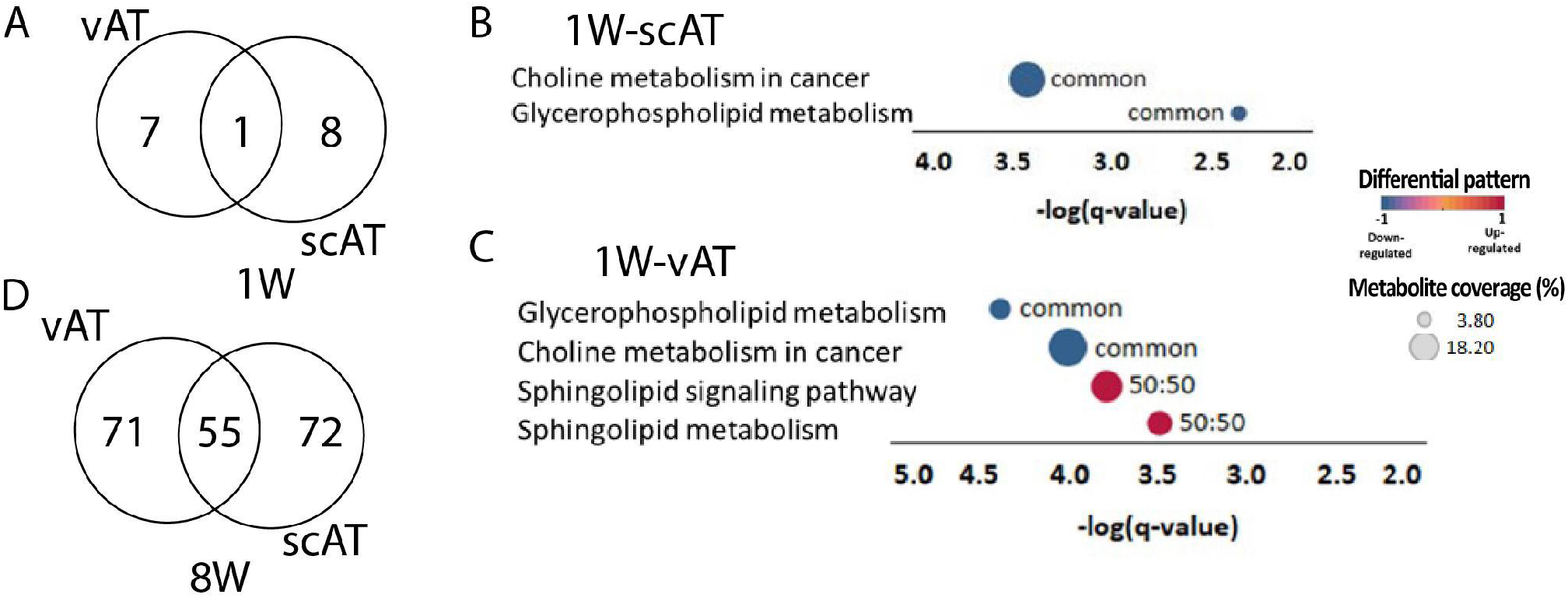
**Changes in the tissue metabolome upon HFD treatment**. The number of differentially abundant metabolites identified in vAT and scAT after 1 week (A) and 8 weeks (D) of HFD treatment. Pathway over-representation analysis was performed in scAT and vAT tissues from mice treated with HFD for 1 week. Significantly enriched pathways (q-value < 0.05) are shown for scAT (B) and vAT (C). Bubble size represents the number of metabolites contributing to each pathway, with larger bubbles indicating a greater number of metabolites. Bubble color represents the direction of metabolite abundance, with red indicating higher abundance of metabolites in HFD-treated mice compared with controls. Unique: >50% of the altered metabolites are specific to the tissue. Common: <50% of the altered metabolites are specific to the tissue. 50:50: 50% of the altered metabolites are specific to the tissue.

**Figure S3.**
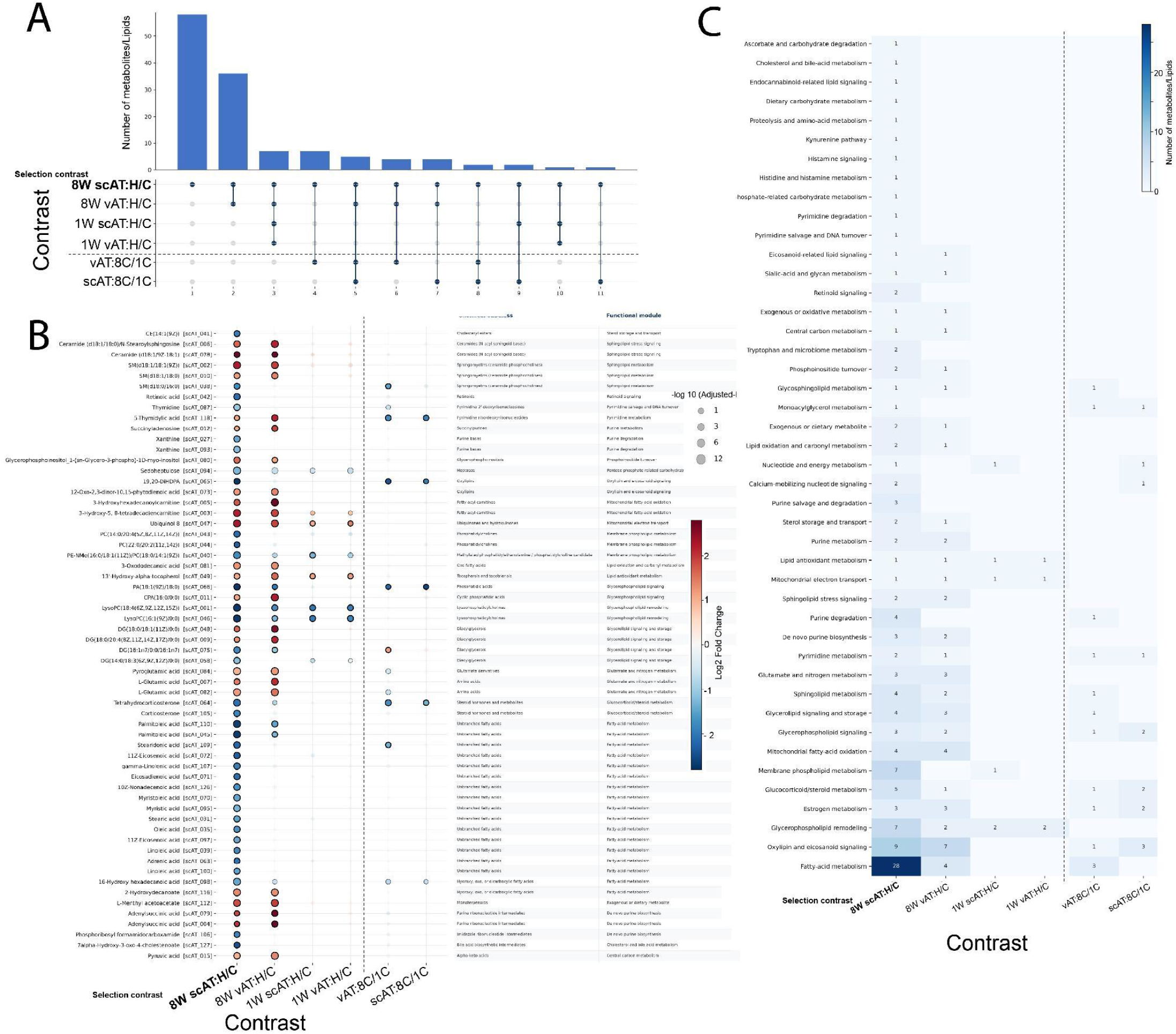
Cross-depot distribution and inferred temporal emergence of the 8-week scAT metabolomic response to HFD. (A) UpSet plot showing intersections of significant features (BH-adjusted P < 0.05 & log2fc => 1) across comparisons. Bars indicate intersection size, connected black dots identify comparisons included in each intersection, and grey dots indicate absence from that comparison. (B) Metabolites significantly altered in subcutaneous adipose tissue (scAT) after 8 weeks of HFD relative to control diet were followed across the corresponding 8-week visceral adipose tissue (vAT) comparison, 1-week scAT and vAT diet comparisons, and 8-versus-1-week control-diet comparisons in both depots. Each row represents an LC–MS/MS feature, retained with its feature identifier when repeated putative metabolite names were present. Red indicates a positive log₂ fold change and blue indicates a negative log₂ fold change relative to the denominator condition. Point size represents −log₁₀BH-adjusted P, applied for visualization. Chemical subclass and functional module annotations are shown to the right. (D) Counts of features satisfying BH-adjusted P< 0.05 and lof2FC=>1, summarized by functional module and comparison; darker shading indicates a larger number of features. scAT, subcutaneous adipose tissue; vAT, visceral adipose tissue; HFD, high-fat diet; C, control diet.

**Figure S4.**
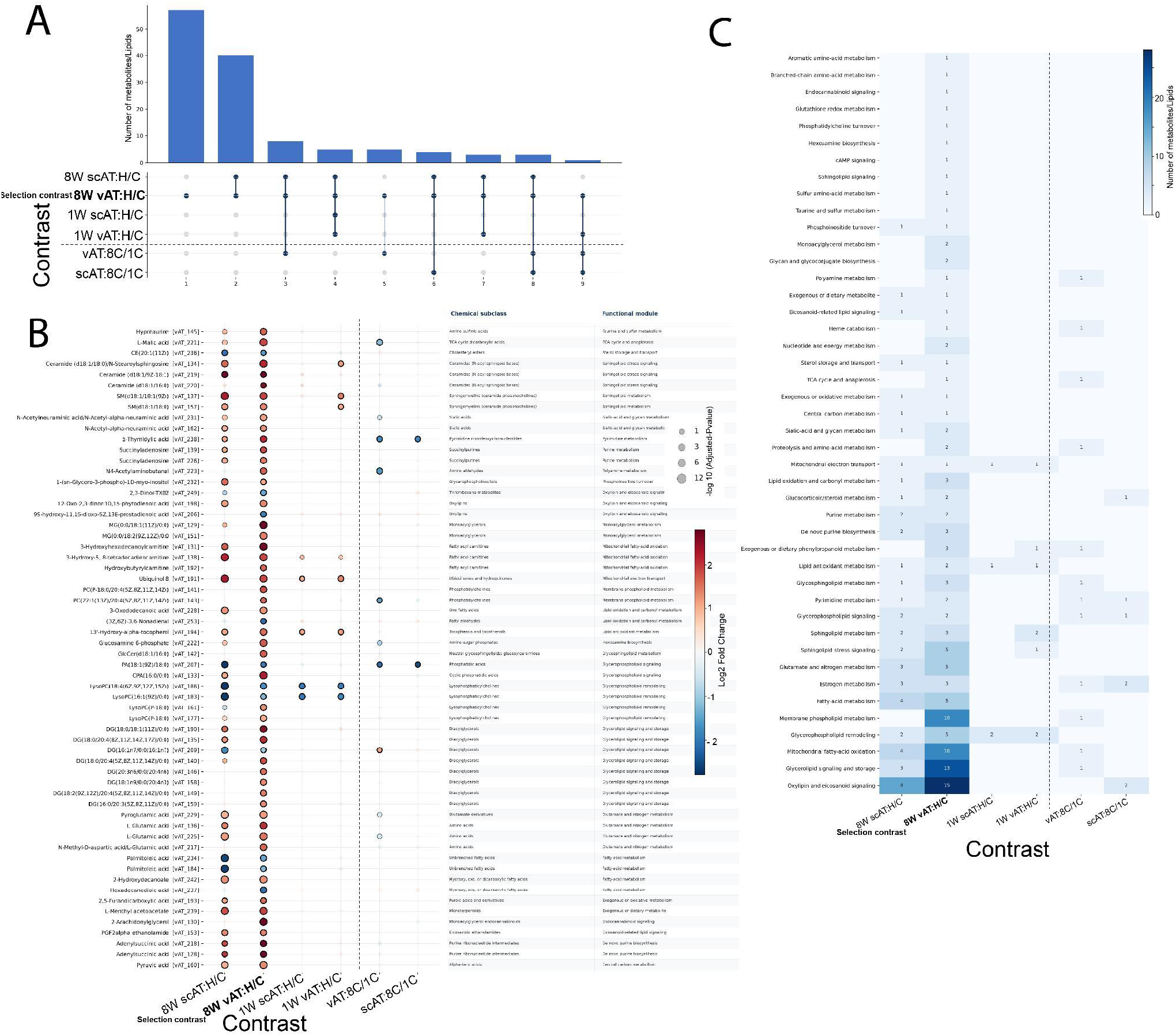
Cross-depot distribution and inferred temporal emergence of the 8-week vAT metabolomic response to HFD. A) UpSet plot showing intersections of adjusted-*P*-significant features across comparisons. Bars indicate intersection size, connected black dots identify comparisons included in each intersection, and grey dots indicate absence from that comparison. (B) Metabolites significantly altered in subcutaneous adipose tissue (vAT) after 8 weeks of HFD relative to control diet were followed across the corresponding 8-week visceral adipose tissue (scAT) comparison, 1-week scAT and vAT diet comparisons, and 8-versus-1-week control-diet comparisons in both depots. Each row represents an LC–MS feature, retained with its feature identifier when repeated putative metabolite names were present. Red indicates a positive log₂ fold change and blue indicates a negative log₂ fold change relative to the denominator condition. Point size represents −log₁₀BH-pvalue, used for visualization. The bigger the size the smaller the BH-adjusted P. Chemical subclass and functional module annotations are shown to the right. (D) Counts of features satisfying BH-adjusted P < 0.05 and lof2FC=>1, summarized by functional module and comparison; darker shading indicates a larger number of features. scAT, subcutaneous adipose tissue; vAT, visceral adipose tissue; HFD, high-fat diet; C, control diet.

**Figure S5.**
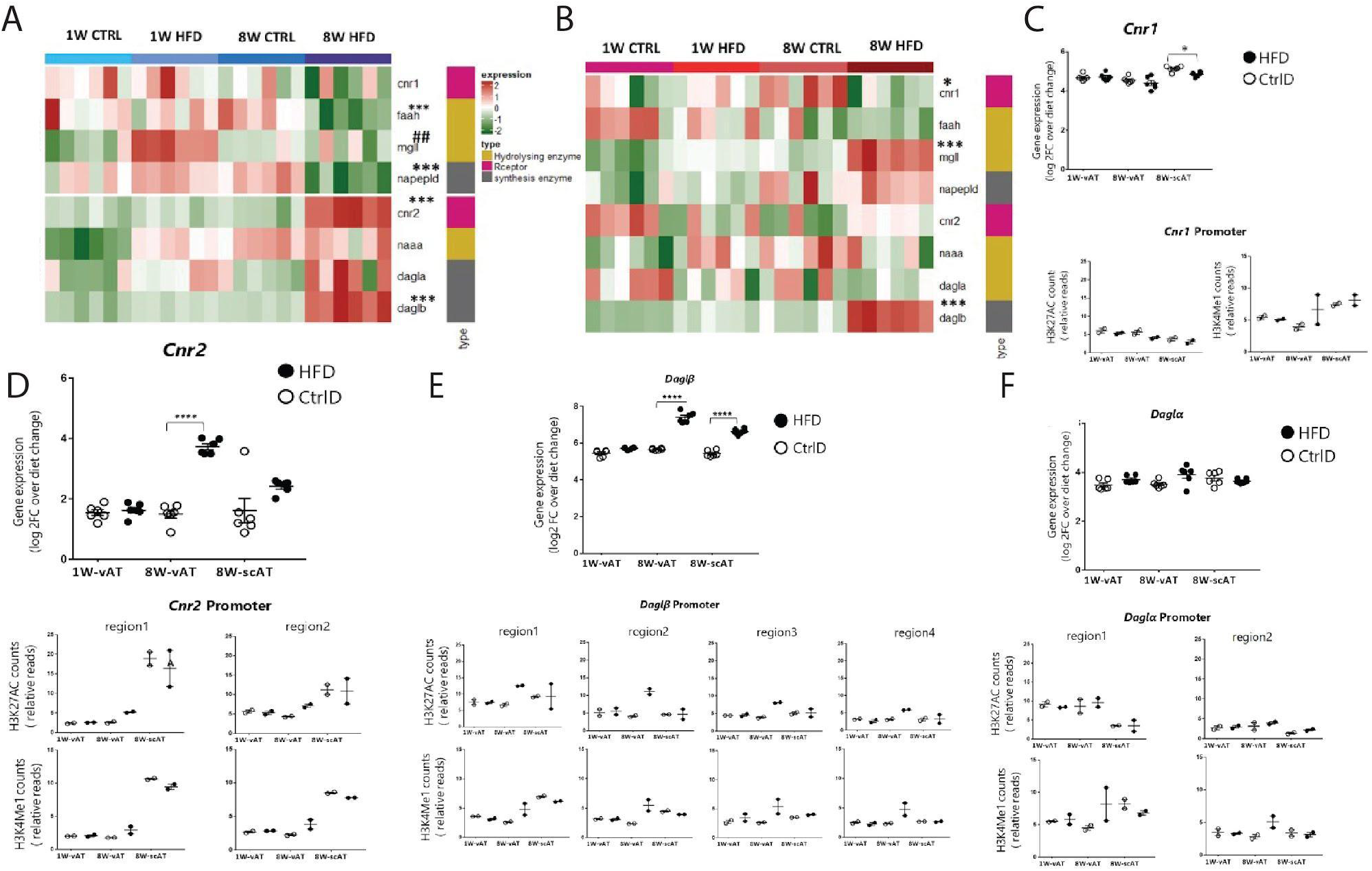
Genes and epigenome landmarks in ECS. A-B) Heatmap of gene expression patterns contributing to ECS in different samples. Impact of obesity driven by HFD on ECS is more straightforward in vAT (A) compared to scAT(B). Two-tailed Student’s t-test was used to calculate the significant changes, for each time point HFD compared to control diet (CtrlD). For 8 week comparison*(p-value < 0.05), ** (p-value < 0.01), *** (p-value < 0.001), ****(p-value < 0.0001) and ## p-value < 0.01 for 1week comparison. C-D) Expression of *cnr1* and *cnr2* in the adipose tissue and its regulation pattern during obesity likewise the profile of histone marks localized at the gene promoters. C) *Cnr1* expression pattern in the vAT and scAT during obesity, comparing HFD-fed with CtrlD-fed mice. Profile of histone marks enriched at the single region in cnr1 promoter demonstrated by H3K27Ac and H3K4Me1 pattern during obesity. (D) *Cnr2* expression profile in the vAT and scAT during obesity in mice received either HFD or CtrlD. (B) Profile of histone marks, H3K27Ac and H3K4Me1, enriched at two regions annotated in *cnr2* promoter during obesity in vAT and scAT. D-E) Enzyme involved in biosynthesis of 2-AG together with epigenetics marks. (D) *Daglβ* expression profile in vAT and scAT during obesity considering two diet models HFD and CtrlD. Histone profile enriched by H3K27Ac and H3K4Me1 at four annotated regions of daglβ promoter during obesity in v-At and scAT. (E) *Daglα* expression profile in vAT and scAT during obesity by feeding mice by HFD or CtrlD. Enrichment patterns of histone marks with virtues of H3K27Ac and H3K4Me1 localized at *Daglα* promoter annotated in two regions during obesity in vAT and scAT. Two-tailed Student’s t-test was used to calculate the significant changes of citrate mean level at different batches compared to the batch one. *(p-value < 0.05), ** (p-value < 0.01), *** (p-value < 0.001), ****(p-value < 0.0001).

**Figure S6:**
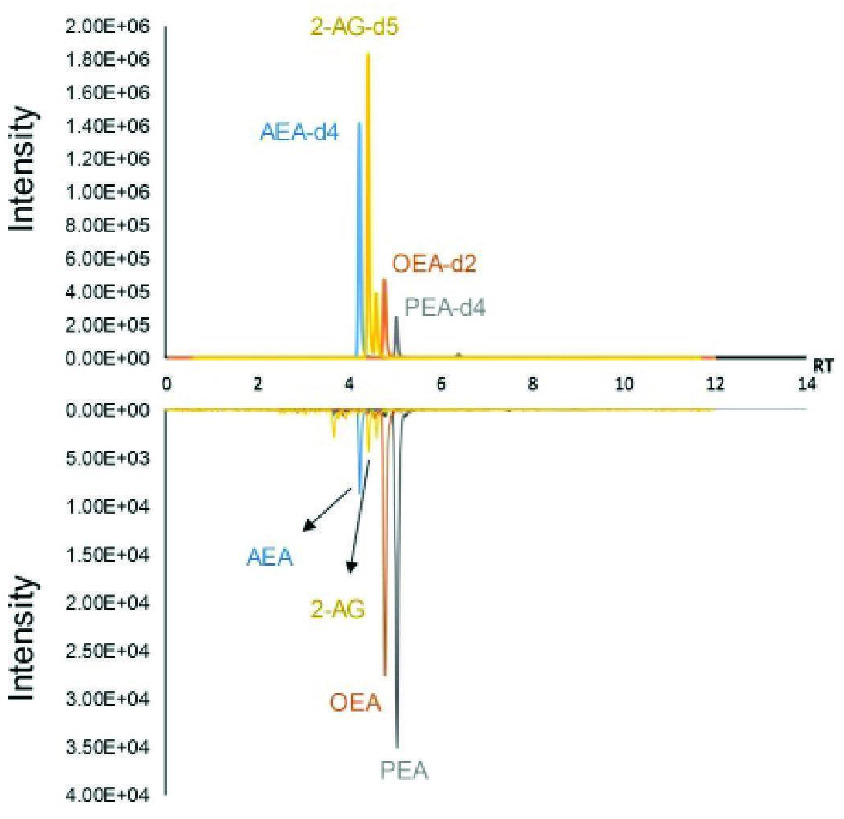
**Representative selected reaction monitoring chromatogram of IS at 400 ng/mL and ECs in adipose tissue’s sample**

**Figure S7:**
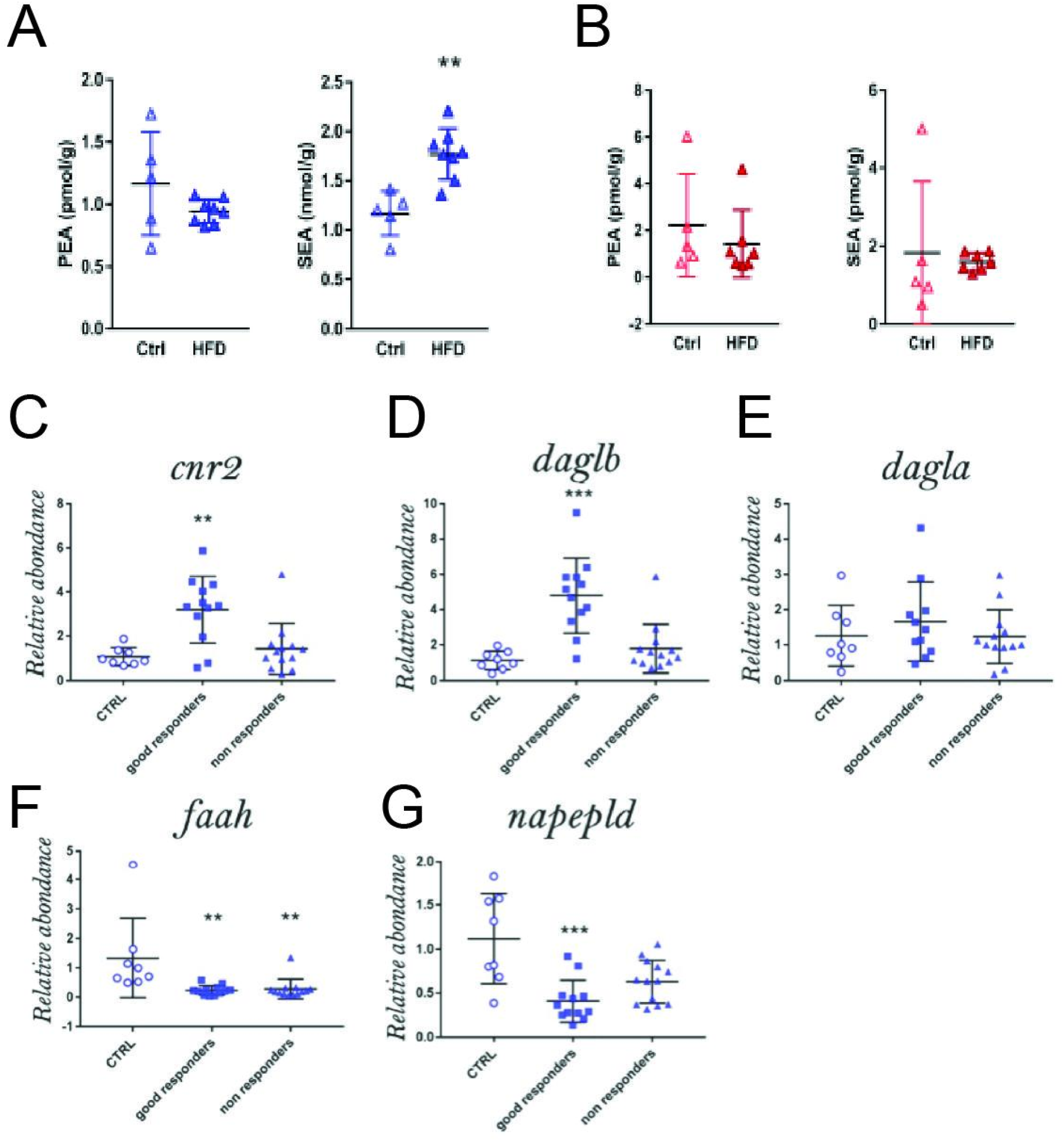
Endocannabinoids coordinated changes. Effect of HFD on the levels of endocannabinoids: Palmitoylethanolamide (PEA) and Stearoylethanolamide (SEA) are measured in vAT (A) and scAT (B). Significant level in comparison with ctrl group has been notified as ** (p-value < 0.01). C-G) Gene expression profile of ECS genes investigated in three experimental groups, chow diet treated mice denoted as Ctrl and HFD treated mice subdivided in good responders (HFD) and non responders, showing low inflammation in vAT (low-INFL HFD). Significant level in comparison with ctrl group has been notified as *(p-value < 0.05), ** (p-value < 0.01), *** (p-value < 0.001).

**Figure S8:**
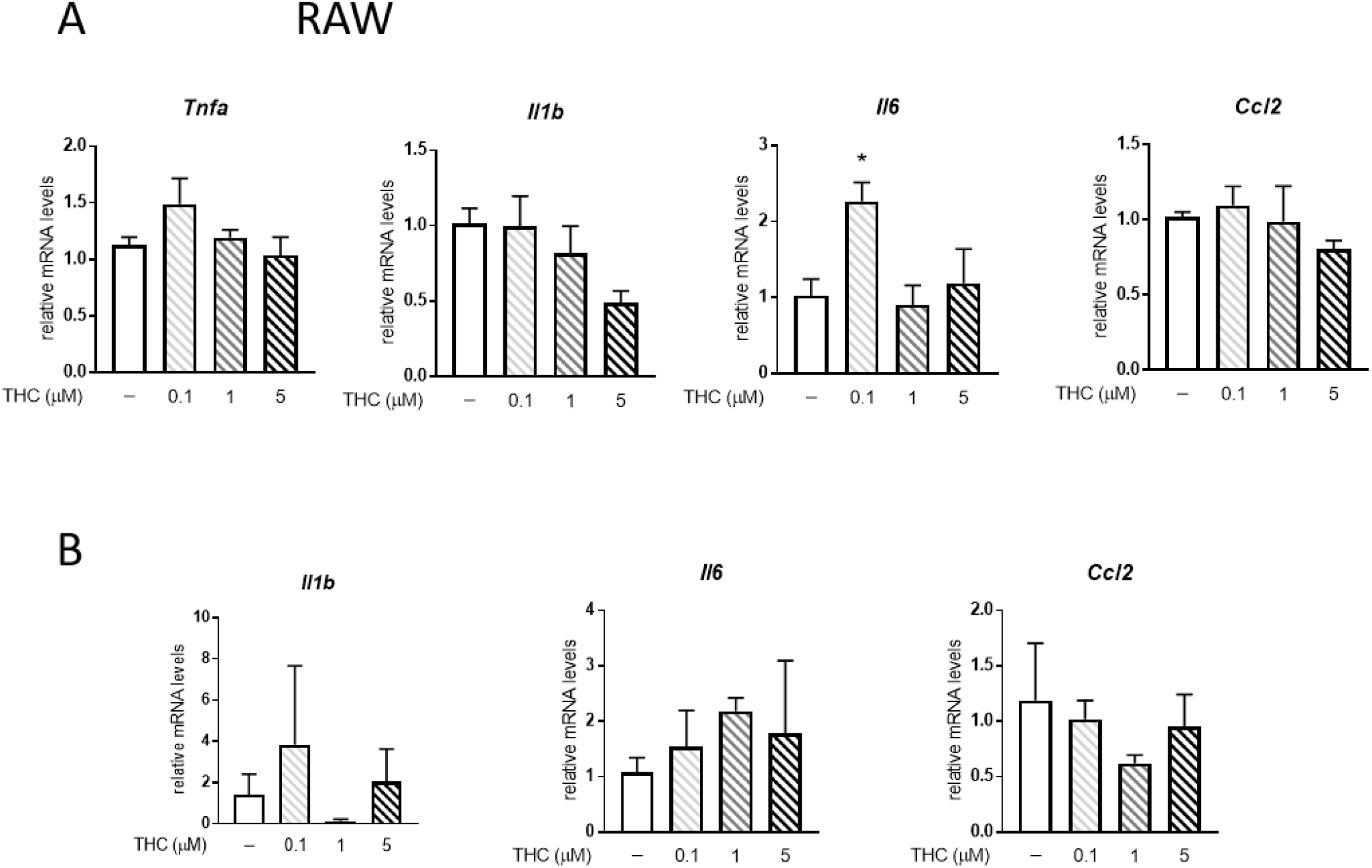
Effect of THC on macrophages and adipocytes. A) RAW264.7 macrophages and B) differentiated 3T3L1 adipocytes were treated with THC at the indicated concentration for 24 hours. mRNA levels of *Tnfa*, *Il1b*, *Il6*, *Ccl2* are presented as mean ± SE (n=3). The significant level of comparative analysis performed between Ctrl (THC -) and treatment groups was *(p-value < 0.05).

**Figure S9.**
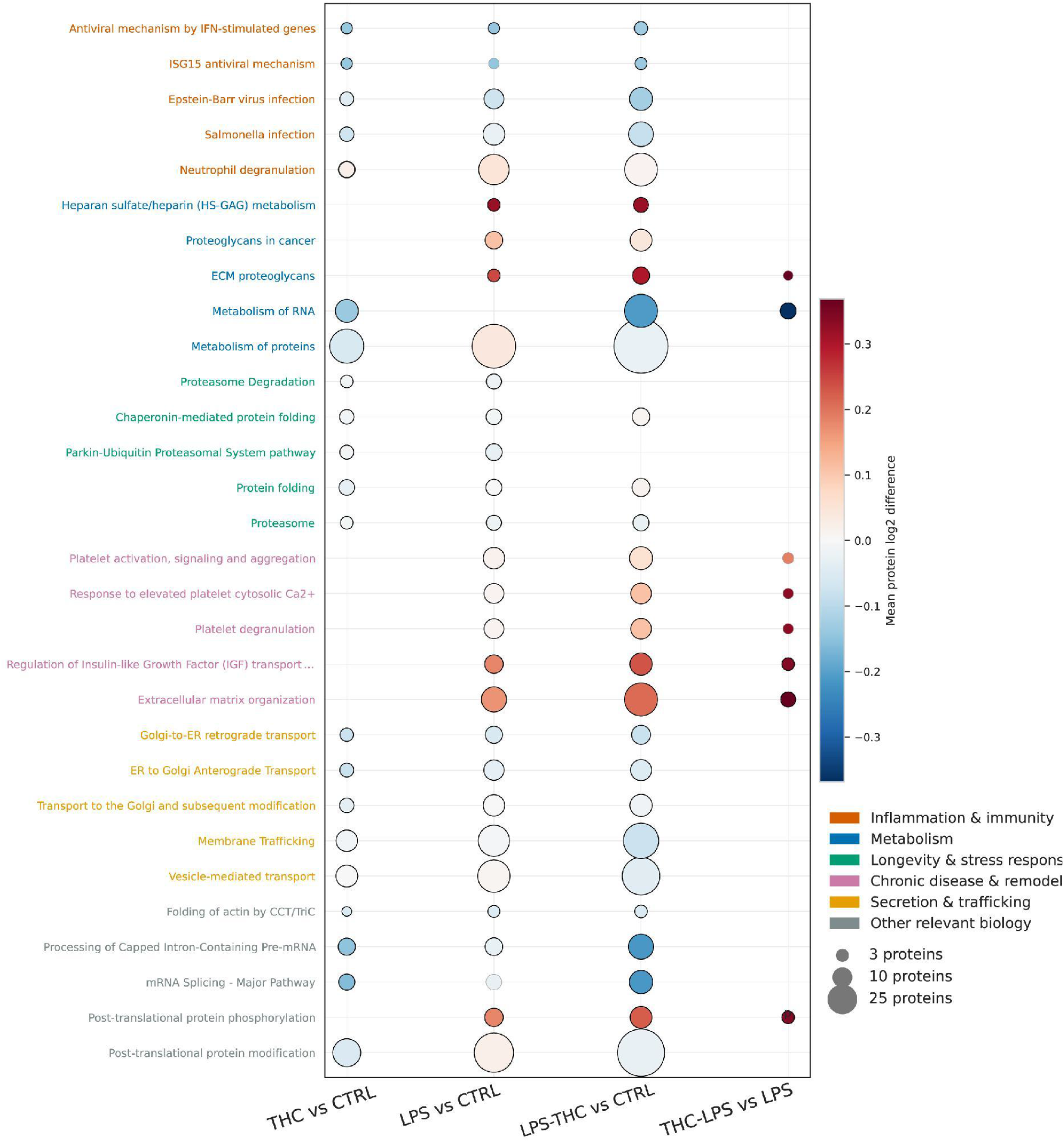
THC remodels the adipocyte-macrophage crosstalk in reponse to LPS. Secreted protein profiling in adipocyte–macrophage co-cultures. Significantly enriched over-representation analysis (ORA) terms were highlighted according to the direction of their mean log₂ fold change. Representative enriched pathways were selected to summarize inflammatory signaling, extracellular-matrix remodeling, metabolism/proteostasis, and secretory trafficking while minimizing redundant pathway annotations.

## Notes

https://iris.unil.ch/entities/publication/423f322b-db50-4370-add5-fd265082903a

### Competing Interest Statement

The authors have declared no competing interest.

## References

1. Hotamisligil, G.S. (2017). Inflammation, metaflammation and immunometabolic disorders. Nature 542, 177–185.

2. Strissel, K.J., Stancheva, Z., Miyoshi, H., Perfield, J.W., 2nd, DeFuria, J., Jick, Z., Greenberg, A.S., and Obin, M.S. (2007). Adipocyte death, adipose tissue remodeling, and obesity complications. Diabetes 56, 2910–2918.

3. Caputo, T., Tran, V.D.T., Bararpour, N., Winkler, C., Aguileta, G., Trang, K.B., Giordano Attianese, G.M.P., Wilson, A., Thomas, A., Pagni, M., et al. (2021). Anti-adipogenic signals at the onset of obesity-related inflammation in white adipose tissue. Cell. Mol. Life Sci. 78, 227–247.

4. Hotamisligil, G.S., Shargill, N.S., and Spiegelman, B.M. (1993). Adipose expression of tumor necrosis factor-alpha: direct role in obesity-linked insulin resistance. Science 259, 87–91.

5. Sugiura, T., Kondo, S., Sukagawa, A., Nakane, S., Shinoda, A., Itoh, K., Yamashita, A., and Waku, K. (1995). 2-Arachidonoylglycerol: a possible endogenous cannabinoid receptor ligand in brain. Biochem. Biophys. Res. Commun. 215, 89–97.

6. Mechoulam, R., Ben-Shabat, S., Hanus, L., Ligumsky, M., Kaminski, N.E., Schatz, A.R., Gopher, A., Almog, S., Martin, B.R., and Compton, D.R. (1995). Identification of an endogenous 2-monoglyceride, present in canine gut, that binds to cannabinoid receptors. Biochem. Pharmacol. 50, 83–90.

7. Côté, M., Matias, I., Lemieux, I., Petrosino, S., Alméras, N., Després, J.-P., and Di Marzo, V. (2007). Circulating endocannabinoid levels, abdominal adiposity and related cardiometabolic risk factors in obese men. Int. J. Obes. (Lond) 31, 692–699.

8. Blüher, M., Engeli, S., Klöting, N., Berndt, J., Fasshauer, M., Bátkai, S., Pacher, P., Schön, M.R., Jordan, J., and Stumvoll, M. (2006). Dysregulation of the peripheral and adipose tissue endocannabinoid system in human abdominal obesity. Diabetes 55, 3053–3060.

9. Sajic, T., Ferreira Gomes, C.K., Gasser, M., Caputo, T., Bararpour, N., Landaluce-Iturriria, E., Augsburger, M., Walter, N., Hainard, A., Lopez-Mejia, I.C., et al. (2024). SMYD3: a new regulator of adipocyte precursor proliferation at the early steps of differentiation. Int. J. Obes. (Lond) 48, 557–566.

10. Bararpour, N., Gilardi, F., Carmeli, C., Sidibe, J., Ivanisevic, J., Caputo, T., Augsburger, M., Grabherr, S., Desvergne, B., Guex, N., et al. (2021). DBnorm as an R package for the comparison and selection of appropriate statistical methods for batch effect correction in metabolomic studies. Sci. Rep. 11, 5657.

11. Kang, S.-C., Kim, B.-R., Lee, S.-Y., and Park, T.-S. (2013). Sphingolipid metabolism and obesity-induced inflammation. Front. Endocrinol. (Lausanne) 4, 67.

12. Tedesco, L., Valerio, A., Dossena, M., Cardile, A., Ragni, M., Pagano, C., Pagotto, U., Carruba, M.O., Vettor, R., and Nisoli, E. (2010). Cannabinoid receptor stimulation impairs mitochondrial biogenesis in mouse white adipose tissue, muscle, and liver: the role of eNOS, p38 MAPK, and AMPK pathways. Diabetes 59, 2826–2836.

13. Kleiner, S., Mepani, R.J., Laznik, D., Ye, L., Jurczak, M.J., Jornayvaz, F.R., Estall, J.L., Chatterjee Bhowmick, D., Shulman, G.I., and Spiegelman, B.M. (2012). Development of insulin resistance in mice lacking PGC-1α in adipose tissues. Proc. Natl. Acad. Sci. U. S. A. 109, 9635–9640.

14. Lo, K.A., and Sun, L. (2013). Turning WAT into BAT: a review on regulators controlling the browning of white adipocytes. Biosci. Rep. 33, e00065.

15. Di Marzo, V., Côté, M., Matias, I., Lemieux, I., Arsenault, B.J., Cartier, A., Piscitelli, F., Petrosino, S., Alméras, N., and Després, J.-P. (2009). Changes in plasma endocannabinoid levels in viscerally obese men following a 1 year lifestyle modification programme and waist circumference reduction: associations with changes in metabolic risk factors. Diabetologia 52, 213–217.

16. D’Eon, T.M., Pierce, K.A., Roix, J.J., Tyler, A., Chen, H., and Teixeira, S.R. (2008). The role of adipocyte insulin resistance in the pathogenesis of obesity-related elevations in endocannabinoids. Diabetes 57, 1262–1268.

17. Camprubí-Rimblas, M., Tantinyà, N., Artigas, A., and Guillamat-Prats, R. (2025). Pharmacological inhibition of the CCL2-CCR2 axis fails to reduce inflammation in a rat model of acute lung injury. Sci. Rep. 15, 31368.

18. Dinarello, C.A. (2011). Blocking interleukin-1β in acute and chronic autoinflammatory diseases. J. Intern. Med. 269, 16–28.

19. Dowling, P., and Clynes, M. (2011). Conditioned media from cell lines: a complementary model to clinical specimens for the discovery of disease-specific biomarkers. Proteomics 11, 794–804.

20. Poschmann, G., Brenig, K., Lenz, T., and Stühler, K. (2021). Comparative secretomics gives access to high confident secretome data: Evaluation of different methods for the determination of Bona fide secreted proteins. Proteomics 21, e2000178.

21. Sato, A., Shimotsuma, A., Miyoshi, T., Takahashi, Y., Funayama, N., Ogino, Y., Hiramoto, A., Wataya, Y., and Kim, H.-S. (2023). Extracellular leakage protein patterns in two types of cancer cell death: Necrosis and apoptosis. ACS Omega 8, 25059–25065.

22. Vartak, R., Deng, J., Fang, H., and Bai, Y. (2015). Redefining the roles of mitochondrial DNA-encoded subunits in respiratory Complex I assembly. Biochim. Biophys. Acta 1852, 1531–1539.

23. Kowalczuk, L., Matet, A., Dor, M., Bararpour, N., Daruich, A., Dirani, A., Behar-Cohen, F., Thomas, A., and Turck, N. (2018). Proteome and metabolome of subretinal fluid in central serous chorioretinopathy and rhegmatogenous retinal detachment: A pilot case study. Transl. Vis. Sci. Technol. 7, 3.

24. Forchelet, D., Béguin, S., Sajic, T., Bararpour, N., Pataky, Z., Frias, M., Grabherr, S., Augsburger, M., Liu, Y., Charnley, M., et al. (2018). Separation of blood microsamples by exploiting sedimentation at the microscale. Sci. Rep. 8, 14101.

25. Thomas, A., Hopfgartner, G., Giroud, C., and Staub, C. (2009). Quantitative and qualitative profiling of endocannabinoids in human plasma using a triple quadrupole linear ion trap mass spectrometer with liquid chromatography. Rapid Commun. Mass Spectrom. 23, 629–638.

26. Montecucco, F., Lenglet, S., Quercioli, A., Burger, F., Thomas, A., Lauer, E., da Silva, A.R., Mach, F., Vuilleumier, N., Bobbioni-Harsch, E., et al. (2015). Gastric bypass in morbid obese patients is associated with reduction in adipose tissue inflammation via N-oleoylethanolamide (OEA)-mediated pathways. Thromb. Haemost. 113, 838–850.

27. Quercioli, A., Montecucco, F., Pataky, Z., Thomas, A., Ambrosio, G., Staub, C., Di Marzo, V., Ratib, O., Mach, F., Golay, A., et al. (2013). Improvement in coronary circulatory function in morbidly obese individuals after gastric bypass-induced weight loss: relation to alterations in endocannabinoids and adipocytokines. Eur. Heart J. 34, 2063–2073.

